# Neutrophils induce effective antibody responses to the pneumococcal conjugate vaccine by inhibiting regulatory T cells

**DOI:** 10.1101/2025.10.28.685204

**Authors:** Essi YI Tchalla, Anagha Betadpur, Andrew Y Khalil, Elena Lopez, Manmeet Bhalla, Musea Chang, Elizabeth A. Wohlfert, Elsa N. Bou Ghanem

**Affiliations:** Department of Microbiology and Immunology, University at Buffalo School of Medicine, Buffalo, NY 14203, USA

**Keywords:** Neutrophils, Pneumococcal conjugate vaccine, *Streptococcus pneumoniae*

## Abstract

Neutrophils are required for production of protective antibodies in response to the pneumococcal conjugate vaccine (PCV), however, the mechanisms behind that are not known. Here, in exploring mechanisms, we found that depletion of neutrophils at the time of vaccination in female mice led to increased regulatory T cells (Tregs) in the spleen that was accompanied by a dysregulated cytokine environment. Following vaccination, neutrophils homed into secondary lymphoid organs where they increased expression of T cell engaging/inhibiting markers. Depletion of Tregs in neutrophil deficient mice restored antibody function against pneumococcal infection. Importantly, PCV vaccination of human female participants altered neutrophil phenotype and *in vitro* coculture of donor neutrophils with PBMCs at one week following vaccination resulted in lower percentage and proliferation of FOXP3^+^ T cells, showing the clinical relevance of this phenotype. This work reveals a new mechanism by which neutrophils ensure protective antibody responses to vaccination by suppressing Tregs.

## INTRODUCTION

Neutrophils or polymorphonuclear cells (PMNs) are innate immune cells that are indispensable for the clearance of many infections and that have been traditionally studied as anti-microbial effector cells (Simmons et al., 2021). However, in the past decade, PMNs have become increasingly appreciated for their ability to orchestrate adaptive immune responses (Bert et al., 2023; Mantovani et al., 2011). In fact, they can affect B and T cells directly or via control of antigen presenting cells such as dendritic cells and macrophages (Leliefeld et al., 2015; Li et al., 2019; Shafqat et al., 2023). They can engage and stimulate T cells in response to antigens via expression of MHC-II, CD80, CD86 and/or CD11c (Abi Abdallah et al., 2011; Culshaw et al., 2008; Matsushima et al., 2013; Vono et al., 2017) or inhibit T cells via production of IL-10, reactive oxygen species, arginase or by expressing PD-L1 (de Kleijn et al., 2013; Kasten et al., 2010; Pillay et al., 2012; Schmielau and Finn, 2001). The ectonucleotidases CD73 and CD39 have been associated with T cell inhibition via adenosine production and are also expressed on PMNs (Allard et al., 2017; Jiang et al., 2023). PMNs can also suppress B cell responses to protein antigens (Bird et al., 2017; Jee et al., 2015; Kamenyeva et al., 2015) or activate B cell responses to bacterial carbohydrates through secretion of B cell activating factor (BAFF), A proliferation-inducing ligand (APRIL), IL-21 or Pentraxin 3 (Chorny et al., 2016; Puga et al., 2011).

While most of the work examining PMN interaction with adaptive immune cells have used model antigens, a few studies have recently focused on the role of PMNs in vaccine responses (Musich et al., 2018). In an adenovirus-SIV vaccination model of Rhesus macaques, PMNs were found to migrate to the vaccine draining lymph nodes (vLNs) and acquire a B cell help phenotype as indicated by expression of BAFF, IL-21 and TNFα among others (Musich et al., 2018). PMNs were found to be required for effective immune responses to both tuberculosis and leishmaniasis live-attenuated vaccine models (Bhattacharya et al., 2020; Trentini et al., 2016). In the context of a clinically relevant vaccine, we showed that PMNs are required at the time of vaccination for induction of protective antibody responses to the licensed pneumococcal conjugate vaccine (PCV) Prevnar (Tchalla et al., 2020). In fact, depleting PMNs at the time of PCV vaccination led to a defect in antibody isotype switching to IgG2c and IgG3, a decline in antibody function, and decreased ability of the mice to control and survive a subsequent pneumococcal infection (Tchalla et al., 2020). However, the mechanisms by which PMNs control antibody responses to PCV remain unclear. This is of high clinical relevance as PCV protects against infections caused by the bacteria *Streptococcus pneumoniae* (pneumococcus), which remain a leading cause of community acquired pneumonia globally, particularly in vulnerable populations such as older adults (CDC, 2022). Understanding the mechanisms behind vaccine efficacy can lead to design of better vaccines.

Conjugate vaccines such as PCV induce T follicular helper cells (Tfh) that aid the production of antibodies by B cells (Rappuoli, 2018; Vinuesa et al., 2016). In fact, PCV vaccination in humans induced the formation of Tfhs, which was linked to enhanced antibody responses (Sterrett et al., 2020). Following vaccination with a conjugate vaccine, germinal centers (GC) form and mature within secondary lymphoid organs such as the lymph nodes (LNs) and spleen (De Silva and Klein, 2015; Pollard et al., 2009). Inside GCs, polysaccharide-specific B cells acquire conjugate antigens from the surface of follicular dendritic cells and process them intracellularly for presentation to Tfhs (De Silva and Klein, 2015; Pollard et al., 2009; Rappuoli et al., 2019a). Interaction between the Tfhs and GC B cells triggers B cell differentiation into antibody secreting cells and enhances antibody production, class switching and affinity maturation (Murphy et al., 2017; Rappuoli et al., 2019b). These antibodies are required for efficient host protection against infections as they facilitate binding, uptake, and killing of bacteria by phagocytes (Forthal, 2014; Vidarsson et al., 2014). Antibody responses by B cells are also highly influenced by cytokines, which are secreted by Tfhs as well as other T cell subsets (Dong, 2021; Vazquez et al., 2015; Vinuesa et al., 2016).

A subset of T cells known to control immune responses are regulatory T cells (Tregs). Tregs have suppressive capacity and play an essential role in homeostasis and immune tolerance (Shevyrev and Tereshchenko, 2019). In the context of vaccination, Tregs have been shown to limit protective immune responses (Bayry, 2014; Moore et al., 2005; Ndure and Flanagan, 2014; Pere et al., 2012; Stepkowski et al., 2024; Weingartner and Golding, 2017). In multiple HIV and cancer models of vaccination, Tregs were shown to dampen both cellular and humoral immune responses, leading to viral replication or tumor progression (Brezar et al., 2015; Togashi et al., 2019). In a pneumococcal polysaccharides model of vaccination, blocking the Treg effector molecule CTLA-4 enhanced antibody and cytokine responses to the polysaccharides (Boudewijns et al., 2005). In humans, higher Tregs in the PBMC of older adults were associated with lower CD4^+^ T cell responses to immunogenic pneumococcal proteins (He et al., 2021). A subset of Tregs that gain access to B cell follicles are termed T follicular regulatory cells or Tfr (Lu and Craft, 2021; Maceiras et al., 2017). Tfrs display phenotypic markers of T follicular helper cells (Tfhs), in addition to their expression of FOPX3 (Maceiras et al., 2017; Qi et al., 2023). They play a role in humoral responses to antigens that is context dependent and have been shown to inhibit B cell responses and accelerate GC contraction (Jacobsen et al., 2021; Sage et al., 2016; Xie et al., 2020). Several studies have looked at the relationship between Tregs and PMNs in models of inflammation and disease and found that Tregs modulate PMN recruitment to the site of inflammation (Himmel et al., 2011; Lewkowicz et al., 2013; Richards et al., 2010). PMNs in turn also promote Treg recruitment in autoimmune and cancer models, and their depletion leads to a decrease in Tregs (Gao et al., 2015; Mishalian et al., 2014; Nadkarni et al., 2016; Perobelli et al., 2016). The interaction of PMNs and Tregs during vaccine responses (including PCV) has not been studied.

In this study, we tested the effect of PMNs on adaptive immune responses to PCV using a preclinical murine model of vaccination as well as by examining responses in vaccinated human participants. We found that following vaccination, PMNs acquire a T cell engaging and suppressing phenotype and suppress Tregs, which is required for the production of protective antibodies. This study reveals a new mechanism by which PMNs shape vaccine responses through the control of Tregs and provides novel targets for the design of more effective vaccines against bacterial infections.

## RESULTS

### PMNs home into secondary lymphoid organs in response to PCV

In prior work, we showed that PMNs are required at the time of PCV administration for the induction of effective antibody response and optimal host protection against pneumococcal disease (Tchalla et al., 2020). There is a well-documented sex-based difference in responses to PCV where females elicit higher antibody responses and vaccine efficacy compared to males (Soneji and Metlay, 2011; Wagenvoort et al., 2017; Wiese et al., 2016) and we recapitulated this in preclinical murine models (Tchalla et al., 2024). Therefore, to understand mechanisms of protection, we focused our studies on female hosts, that mount higher humoral and cellular responses to PCV and are better protected against subsequent pneumococcal infection compared to male hosts (Tchalla et al., 2024).

To better understand how PMNs control PCV efficacy, we first asked whether they are recruited to secondary lymphoid organs (spleen and vaccine draining lymph nodes (vLNs)) in response to vaccination. We looked at expression of migration markers CXCR4 and CCR7 on circulating PMNs 24 hours post vaccination as they are involved in PMN trafficking to the lymph nodes via the lymphatic system or high endothelial venules (Fig. S1 & Fig. 1A) (Beauvillain et al., 2011; Hampton et al., 2015). We found that at baseline, a low percentage of circulating PMNs express CXCR4 and CCR7 and upon vaccination, CCR7^+^ PMNs significantly increase (Fig. 1A). This higher percentage of CCR7^+^ PMNs coincided with a decrease in the number of circulating PMNs 18 to 24 hours post vaccination and an influx of PMNs to the spleen and vLNs at those time points respectively (Fig. 1B). Following the peak of PMN influx into the spleen and vLNs, PMN numbers return to baseline by 48 hours post vaccination (Fig. 1B). These findings demonstrate that PMNs are rapidly recruited to secondary lymphoid organs in response to PCV.

**Figure 1:**
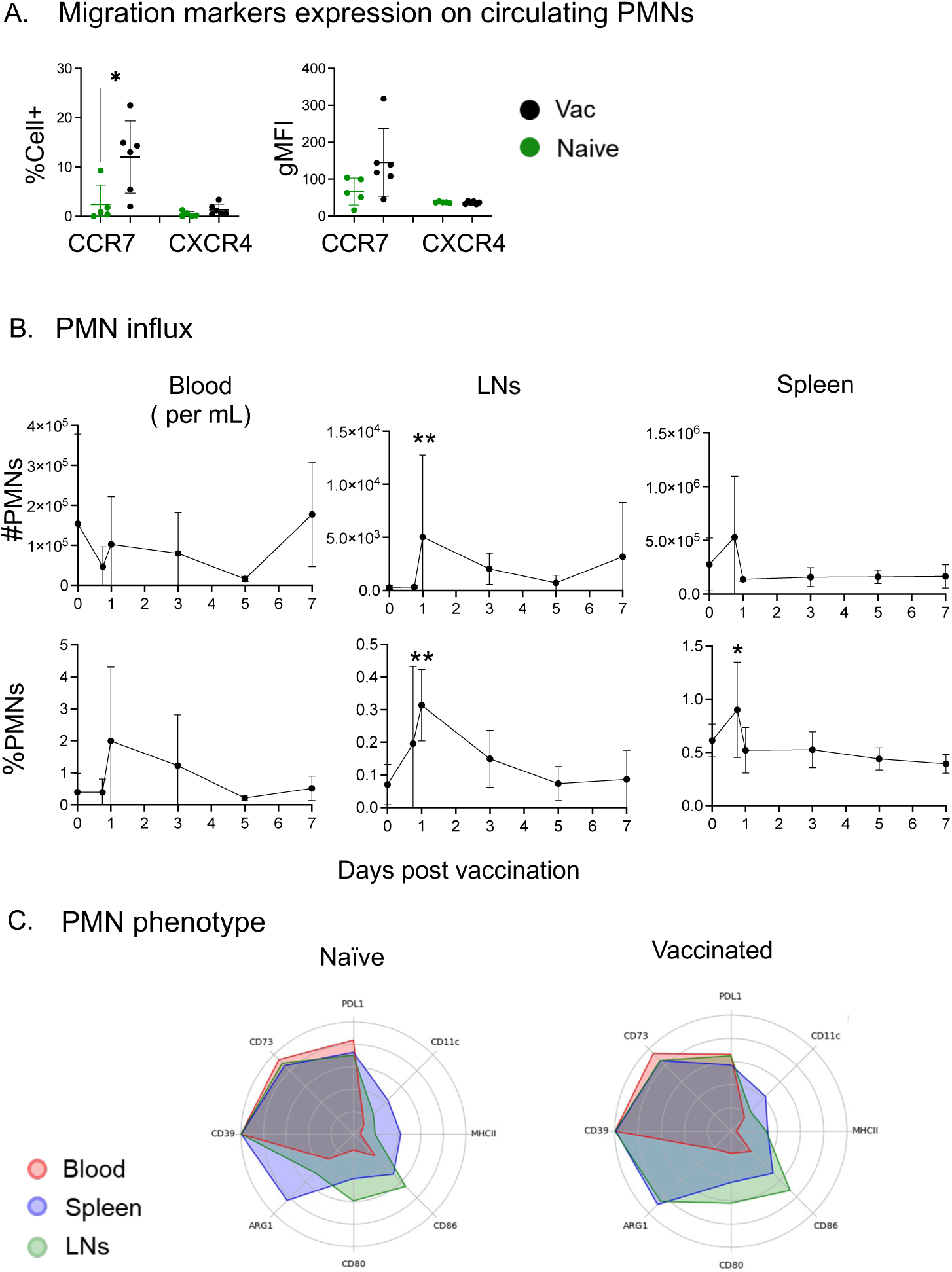
PMN migration to secondary lymphoid organs following vaccination. Adult C57BL/6 female mice were injected with PCV or PBS. (A) Blood was collected from the mice 24 hours following vaccination and assessed using flow cytometry. Percentages (left) and gMFI (right) of CCR7 and CXCR4 on live PMNs (Ly6G^+^CD11b^+^) were determined. (B) At different timepoints post vaccination, blood, spleen and vLNs were collected from all mice and live PMN (Ly6G^+^CD11b^+^) numbers (top) and percentages (bottom) was assessed using flow cytometry. (C) Post vaccination, blood, spleen and vLNs were collected and percent expression of various markers on PMNs were assessed by flow cytometry. Expression of MHC-II PD-L1, ARG1, CD73 and CD39, CD80, CD86 and CD11c were assessed 18-24 hours post vaccination. Data show percent expression of markers on PMNs in all organs/tissue overlayed for naïve and vaccinated mice. (A) Data are pooled from 3 separate experiments and each dot represents an individual mouse with a total of n=5 naïve and n=6 vaccinated mice. Graphs represent the mean +/− SD. * denotes significant differences between the indicated groups as determined by One-way ANOVA followed by Sidak’s multiple comparisons test. * is *p*≤0.05. (B) Data are pooled from 3 separate experiments with a total of 6 mice per timepoint. Line graphs represent the mean +/− 95% confidence interval. * denotes significant differences between the indicated groups as determined by Kruskal-Wallis test followed by Dunn’s multiple comparisons test. * is *p*≤0.05 and ** is *p*≤0.01. (C) Data are pooled from 3 separate experiments with a total of n=5 naïve and n=6 vaccinated mice and are presented as radar plots generated using R studio. Abbreviations: gMFI = geometric Mean Fluorescence Intensity, MHC = Major Histocompatibility Complex.

### PMNs do not affect B cell numbers in response to PCV

To start investigating how PMNs induce protective antibodies to PCV, we first examined changes to the B cell compartment. We depleted PMNs at the time of vaccination as presented in Fig S2A and previously described (Tchalla et al., 2020). The efficiency of PMN depletion was assessed over 28 days post vaccination in the spleen (Fig. S2B). We observed >90% PMN depletion at day 7 (Fig. S2B) with a return to steady state numbers in the spleen by day 28 (Fig. S2B) (Tchalla et al., 2020). We then investigated B cell responses post PMN depletion in the spleen on days 7, 14 and 21 post vaccination (Fig. S3A). Upon PMN depletion, there was no change in the percentage and numbers of total B cells (B220^+^) in the spleen compared to isotype treated mice at any of the timepoints investigated (Fig. S4A). Similarly, no effect of PMN depletion was observed on GC B cells (CD38^lo^CD96^+^) and plasmablasts (CD138^+^) (Fig. S4B & S4C). In prior work, we had observed a defect in isotype switching, opsonic and binding capacities of antibodies in response to PCV in PMN deficient hosts (Tchalla et al., 2020), indicating a possible defect on somatic hypermutation (SHM) and/or class switch recombination (CSR). Therefore, on days 7 post vaccination, we assessed B cell expression of Activation Induced Cytidine Deaminase (AID), an enzyme required for both SHM and CSR (Di Noia and Neuberger, 2007; Stavnezer et al., 2008) (Fig. S3B). For both total and GC B cells, there was an increase in AID expression in mice following vaccination (Fig. S5). However, depleting PMNs had no effect on the percentage of AID^+^ B cells or the expression of AID by B cells (Fig. S5). These findings suggest that PMNs are not directly acting on B cells during vaccination.

### PMNs do not affect macrophages and dendritic cells in response to PCV

We then looked at the effect of PMNs on other immune cells important for vaccine responses such as antigen presenting cells (APCs) (Abi Abdallah et al., 2011; Morel et al., 2008). For that, we depleted PMNs at the time of vaccination (Fig. S2) and analyzed the percentage of splenic macrophages (F4/80^+^) and dendritic cells (DC-CD11c^+^) or their respective expression of antigen presenting molecules MHC-II, CD80 and CD86 on days 7, 14 and 21 post vaccination (Fig. S6). Overall, we found that depletion of PMNs had no effect on the percentages of both macrophages and DCs present in the spleen (Fig. S7). No substantial difference in expression of CD80, CD86 and MHC-II by DCs and macrophages was observed between PMN depleted and PMN sufficient mice (Fig. S7). These findings suggest that PMNs are not directly acting on macrophages or DCs immediately upon vaccination.

### Absence of PMNs at the time of vaccination alters the cytokine environment in the spleen

The cytokine environment in secondary lymphoid organs controls the quality of antibodies produced (Abbas et al., 1993). Therefore, we next looked more broadly at the overall inflammatory environment in the spleen in the presence or absence of PMNs in vaccinated mice using a cytokine protein array. We found that overall, in the absence of PMNs at the time of vaccination, there was an increase in inflammatory mediators in the splenic milieu compared to when PMNs are present (Fig. S8). This increase in inflammation was seen at day 7 and resolved by day 14 through 21 post vaccination (Fig. S8), timepoints that coincide with return of PMNs after depletion (Fig. S2B) (Tchalla et al., 2020). These data show that the absence of PMNs at the time of vaccination leads to an altered inflammatory response in the spleen, suggesting a regulatory role for PMNs during PCV.

We had observed a defect in isotype switching to IgG2 and IgG3 antibodies in the absence of PMNs (Tchalla et al., 2020), therefore, we directly tested whether PMNs controlled effective antibody response to PCV through IFNγ, as this cytokine is required for class switching to those isotypes (Karachunski et al., 2000; Murphy et al., 2017). We measured the percentage of IFNγ^+^ PMNs in the spleen 18 hours post vaccination following *ex vivo* stimulation with PMA, heat-killed *S. pneumoniae* or PCV (Fig. S9). We found that more PMNs express IFNγ when stimulated with PMA, however, there was no change in the percentage of IFNγ^+^ PMNs following vaccination and upon restimulation (Fig. S10A). Similarly, total IFNγ levels in the spleen within the first 28 days of vaccination were comparable between PMN-depleted and PMN-sufficient mice (Fig. S10B). Additionally, when we blocked IFNγ *in vivo* at the time of vaccination using an αIFNγ antibody following the same schedule as for the depletion of PMNs shown in Fig. S2A, there was no effect on bacterial burden in the lung and blood at 24 hours post infection (Fig. S10C) or on host survival (Fig. S10D) compared to the isotype-treated mice. To verify that the blocking antibody was functioning, we measured isotype switching to IgG2 which is controlled by IFNγ (Murphy et al., 2017), and founded that blocking IFNγ led to an increase in IgG1 antibodies and decrease in IgG2b and IgG2c (Fig. S10E). In summary, these data indicate that overall, the control of pneumococcal antibody responses by PMNs is not mediated by IFNγ.

### PMNs acquire a T cell regulating phenotype upon entry into secondary lymphoid organs

To better understand how PMNs control PCV efficacy, we assessed how they respond to vaccination. PMNs have been shown to acquire different phenotypes depending on entry into organs/tissues and the composition of the tissue microenvironment (Ballesteros et al., 2020; Yang et al., 2019). Therefore, we examined PMN phenotype within the first 24 hours post vaccination when they influx into secondary lymphoid organs (Fig. 1B). As the cytokine responses in the spleen suggested that PMNs play a regulatory role, we looked at the expression of various T cell engaging (MHC-II, CD80, CD86 and CD11c) and suppressing (PDL-1, CD73, CD39 and ARG1) markers on PMNs (Fig. S1B). In all sites investigated, overall, expression of both T cell engaging and suppressing markers on PMNs did not change upon vaccination except for a decrease in PD-L1^+^ PMNs in the blood and spleen (Fig. S11 & S12). However, there were tissue specific differences regardless of vaccination status (Fig. S11, S12 & 1C). Expression of CD80, CD86, CD11c MHC-II and ARG1 were higher on PMNs in secondary lymphoid organs compared to circulating PMNs in both naïve and vaccinated mice (Fig. S11, S12 & 1C). These data are consistent with the literature showing that entry into tissues instructs PMNs to acquire organ specific phenotypes (Ballesteros et al., 2020; Yang et al., 2019). Additionally, the data suggest that as PMNs influx into secondary lymphoid organs following vaccination (Fig. 1B), they acquire a tissue specific regulatory phenotype. As this PMN population decreases in the blood and enters secondary lymphoid organs (Fig 1B), they may play a role on T cell responses at those sites.

### PMNs do not affect T helper cell responses to PCV

To test the effect of PMNs on T cell responses, we depleted PMNs at the time of vaccination (Fig S2) and measured the percentages and numbers of T helper 1 (Th1) (CD4^+^TCRβ^+^FOXP3^-^Tbet^+^), Th2 (CD4^+^TCRβ^+^FOXP3^-^GATA3^+^), Th17 (CD4^+^TCRβ^+^FOXP3^-^RORγt^+^) and Tfh (CD4^+^TCRβ^+^CXCR5^+^PD1^+^) cells and in the spleen 7-, 14– and 21-days post vaccination (Fig. S13). We found that there was no difference in the percentages or numbers of any of the T cell subsets between vaccinated PMN-depleted and PMN-sufficient mice at any the timepoints investigated in the spleen (Fig. S14). As examining total CD4^+^ T cells may mask changes within the smaller antigen experienced population, we also looked at antigen experienced cells by gating on CD49d^+^CD11a^+^ T cells (Fig. S13A) (Christiaansen et al., 2017). Similarly, we found no effect of PMN depletion on antigen experienced Th1, Th2, Th17 and Tfhs in the spleen (Fig. S15).

As depleting PMNs had no effect on the amount of T helper cells (Fig. S14 & S15), we asked if their ability to produce cytokines was impaired. We examined cytokine responses *ex vivo* using PMA/Ionomycin as positive stimulant for cytokine production. Using flow cytometry, we measured expression of IFNγ, IL-4, IL-17, TGFβ and IL-10 by splenic CD44^hi^ activated CD4^+^ T cells at day 14 post vaccination (Fig. S16). We used the protein Latency Associated Peptide (LAP) as a proxy for TGFβ production since it acts as a chaperone for TGFβ and their levels correlate (Areström et al., 2012; Liu et al., 2021). The cytokine response was skewed towards production of IFNγ and LAP, both in the presence and absence of PMNs, as the percentage of CD4^+^TCRβ^+^ T cells expressing these cytokines was higher than the other cytokines investigated (Fig. S17). Overall, PMA/Ionomycin stimulation increased LAP, IFNγ and IL-4 expression by CD44^hi^ CD4^+^ T cells above the no stimulation condition (Fig. S17A-C). However, PMN depletion did not affect expression of any of the cytokines by T cells in comparison to PMN-sufficient mice (Fig. S17).

### PMNs inhibit Treg expansion in the spleen in response to PCV

We next assessed the effect of PMNs on regulatory T cells (Tregs) (CD4^+^TCRβ^+^FOXP3^+^) and T follicular regulatory cells (Tfrs) (CD4^+^TCRβ^+^CXCR5^+^PD1^+^FOXP3^+^) following PCV (Fig. 2). We saw an increase in the percentage of Tregs in the spleens of PMN-depleted mice compared to PMN-sufficient mice at day 14 post vaccination (Fig. 2A). In PMN-depleted mice, there was also an increase in the percentage and numbers of Tfrs at day 7 as well as an increase in percentage at day 21 post vaccination (Fig. 2B). When looking at the antigen experienced compartment, there was an increase in Tregs at days 7 and 14 post vaccination in the spleen of PMN-depleted mice compared to PMN-sufficient mice (Fig. 2C) and differences in the Tfrs compartment at days 7 and 21 (Fig. 2D). No differences in Tregs or Tfrs were observed in the lymph nodes (data not shown). These data suggest that PMNs inhibit the expansion of regulatory T cells in the spleen following PCV.

**Figure 2:**
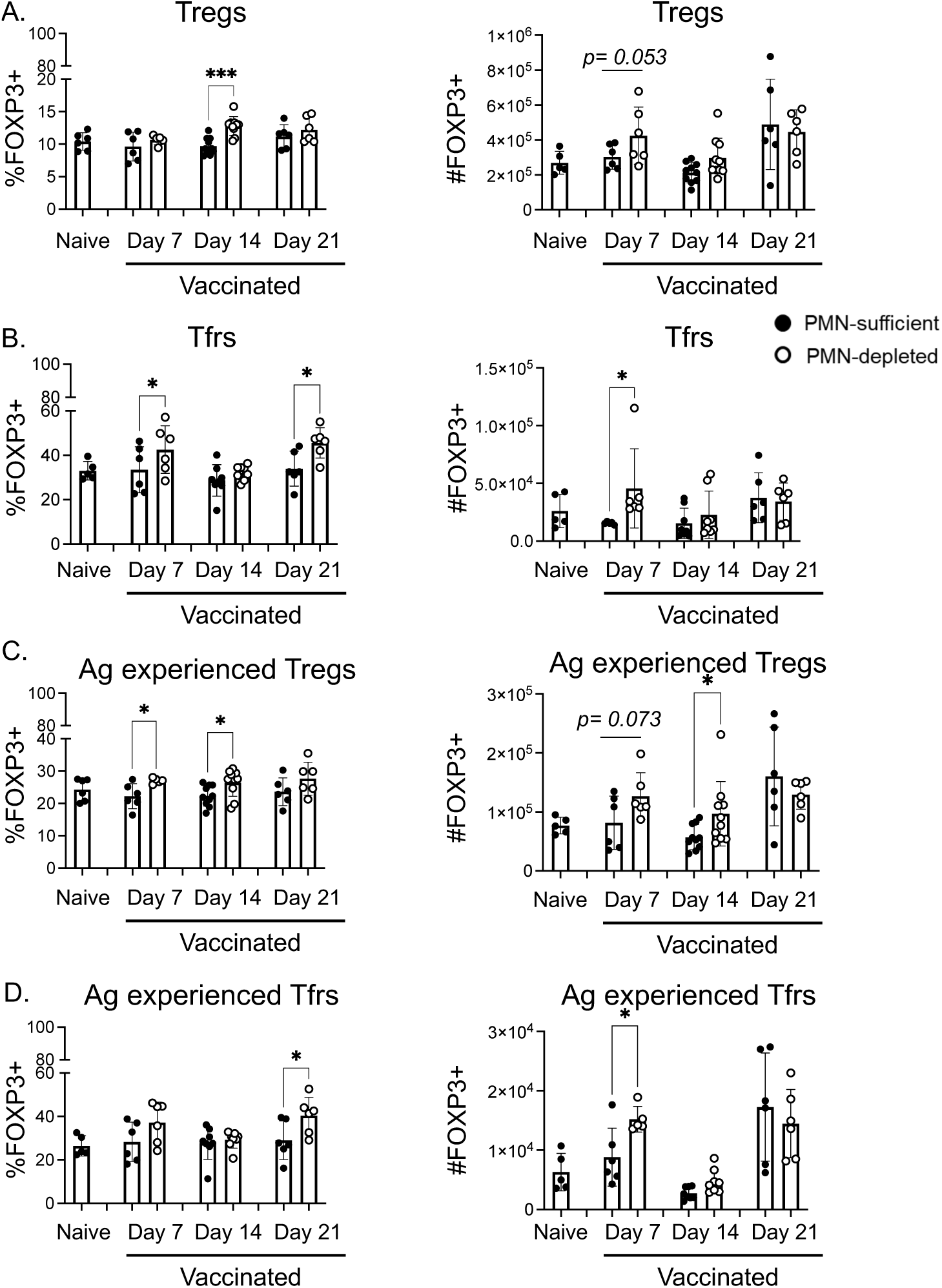
Effect of PMN depletion on regulatory T cells. Adult C57BL/6 female mice were injected with PCV or PBS and depleted of their PMNs or treated with an isotype control. Spleens were collected from all mice and assessed using flow cytometry on days 7, 14 and 21 post vaccination. (A-B) Live total (CD4^+^TCRβ^+^) and (C-D) antigen experienced (CD11a^+^CD49d^+^) T cells were gated on. Percentages (left) and numbers (right) of (A&C) regulatory T cells (FOXP3^+^) and (B&D) T follicular regulatory T cells (CXCR5^+^PD1^+^ FOXP3^+^) were determined from the total and antigen experienced T cell gates. Data are pooled from 4 separate experiments and each dot represents an individual mouse with a total of n=6 naïve, n=6 vaccinated day 7, n=8-10 vaccinated day 14, and n=6 vaccinated day 21 mice per group. * denotes significant differences between the indicated groups as determined by One-way ANOVA followed by Sidak’s multiple comparisons test. * is *p*≤0.05 and *** is *p*≤0.001. Bar graphs represent the mean +/− SD.

### PMNs inhibit Treg expansion in response to PCV *in vitro*

To directly assess the effect of PMNs on Tregs, we used *in vitro* assays. For that, we performed coculture assays of splenocytes isolated from day 1-vaccinated mice in the presence or absence of PMNs using heat-killed bacteria that contain bacterial polysaccharides or αCD3 as stimulants for 48 hours. PMNs were removed from splenocytes using positive isolation magnetic sorting. Similar to what we observed *in vivo*, in the absence of PMNs, there was an increase in the percentage of Tregs in response to stimulation with heat-killed *S. pneumoniae* (Fig. 3). PMN removal did not alter other T helper cell subsets (Fig. S18). This recapitulates the *in vivo* findings and suggests that PMNs control Tregs response to bacterial polysaccharides.

**Figure 3:**
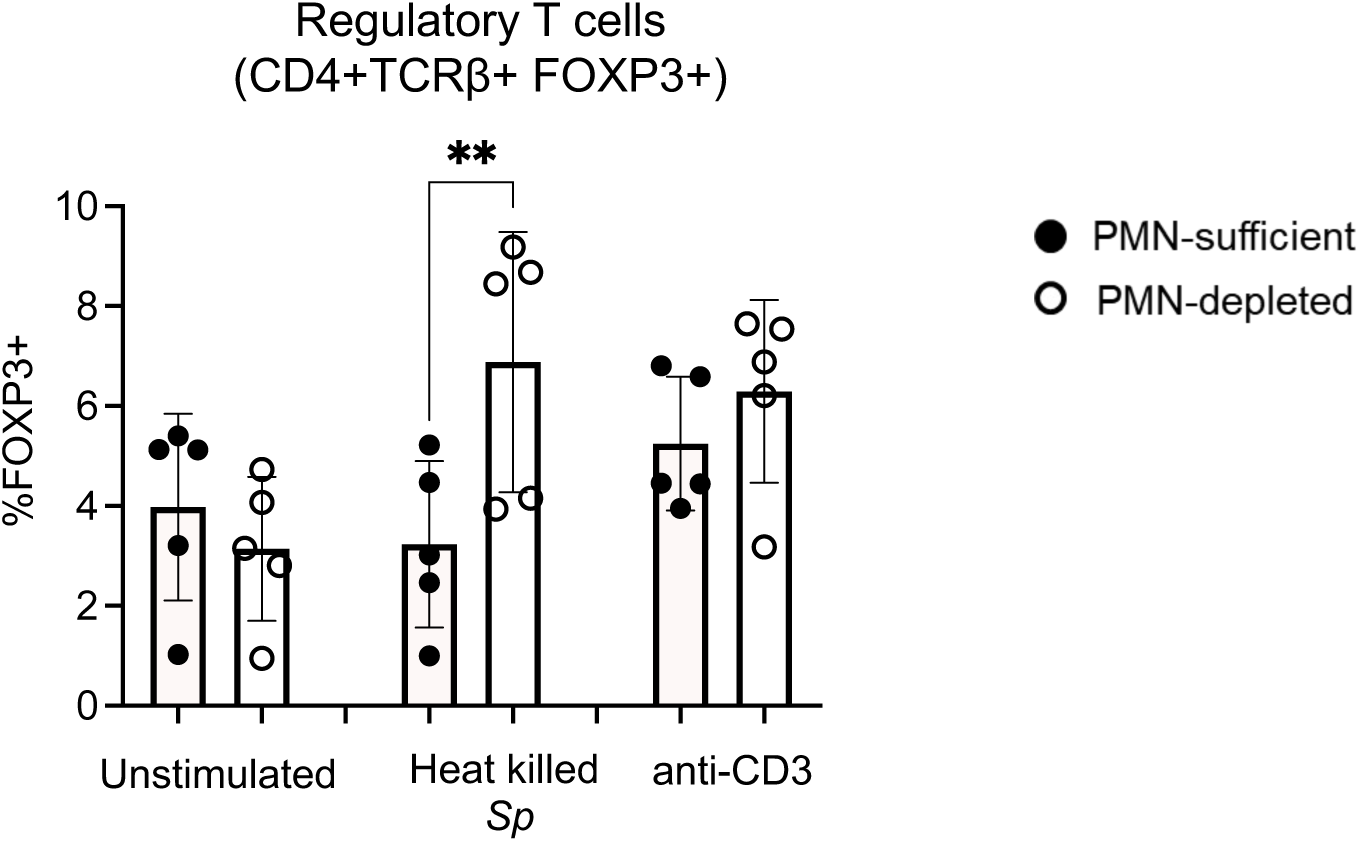
Effect of PMN depletion on Tregs in mice *in vitro*. Adult C57BL/6 female mice were injected with PCV or PBS and depleted of their PMNs or treated with an isotype control. On day 1 post vaccination, splenocytes were isolated and cultured in the presence or absence of PMNs for 2 days with heat-killed *S. pneumoniae* or αCD3 as stimulants. Percentage of FOXP3^+^ cells were determined in the live CD4^+^ cells gate is shown. Data are pooled from separate experiments. * denotes significant differences between the indicated groups as determined by One-Way ANOVA followed by Šídák’s multiple comparisons test. ** is *p*≤0.01. Graphs represent the mean +/− SD.

### Reducing Tregs in PMN-depleted hosts rescues antibody responses

Next, we tested whether the increase in regulatory T cells is responsible for the defective antibody responses seen in the absence of PMNs at the time of vaccination (Tchalla et al., 2020). To investigate this, we depleted Tregs using an αCD25 antibody (Farkas et al., 2013) either alone or in conjunction with PMN depletion using αLy6G (Fig. S19A). We achieved ∼90% depletion of PMNs in the blood and ∼40-50% depletion of Tregs in both spleen and blood (Fig. S19B & S19C). Moreover, αCD25 and αLy6G injections had no effect on percentages of Th1, Th2, Th17, Tfhs and B cells in the spleen (Fig. S19D).

We then followed antibody levels upon Treg and/or PMN depletion in the blood of mice at weeks 0, 2 & 4 post vaccination. We found an increase in total IgG against pneumococcal polysaccharides in PMN-& Treg-depleted mice compared to PMN-depleted mice at 4 weeks post vaccination (Fig. 4A). We next investigated the ability of antibodies in the sera to bind the surface of the pneumococcus (Fig. S20). In all mouse groups, vaccination significantly increased binding of IgG to the bacteria compared to baseline (Fig. 4B). As previously reported, sera from vaccinated PMN-depleted mice had significantly lower IgG binding compared to sera from vaccinated PMN-sufficient mice (Fig. 4B) (Tchalla et al., 2020). Interestingly, depletion of Tregs in PMN-depleted mice rescued IgG binding in the sera to similar levels to that of PMN-sufficient vaccinated mice, while depletion of Tregs alone did not affect IgG binding in the sera of PMN-sufficient vaccinated mice (Fig. 4B). We also looked at the ability of antibodies in the sera to promote opsonophagocytic killing of the pneumococcus by PMNs isolated from naïve mice. As expected, there was an increase in bacterial killing by PMNs when sera from vaccinated versus naïve mice were used as opsonins (Fig. 4C). Similar to above, depletion of Tregs in PMN-depleted mice rescued the ability of the sera to induce opsonophagocytic bacterial killing to levels similar to PMN-sufficient vaccinated mice, while depletion of Tregs alone did not affect the opsonic capacity of sera from PMN-sufficient vaccinated mice (Fig. 4C).

**Figure 4:**
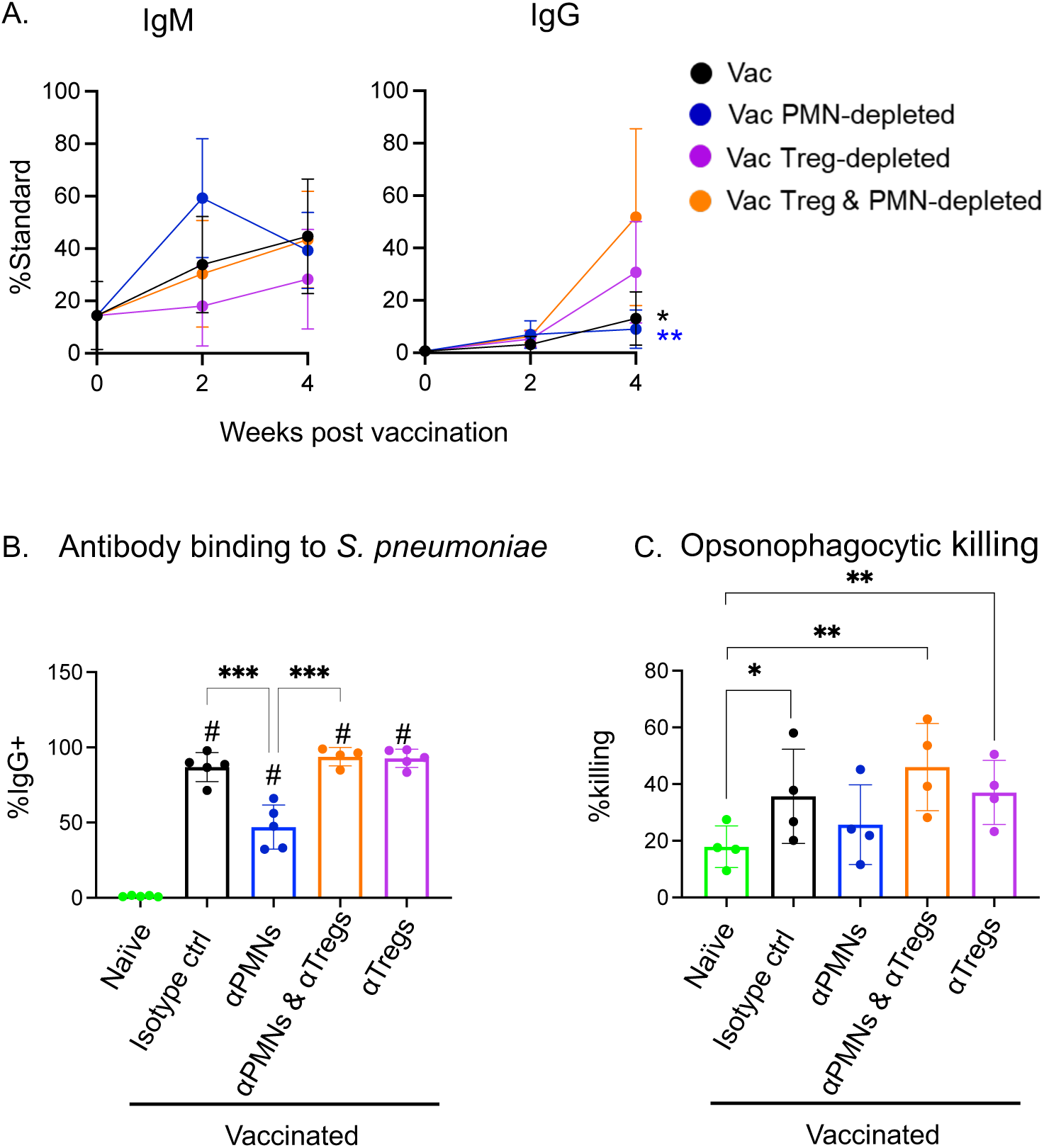
Effect of Treg depletion on antibody responses. Adult C57BL/6 female mice were injected with PCV or PBS and depleted of their PMNs and/or Tregs (αCD25) or treated with an isotype control. (A) Production of IgM and total IgG against type 4 pneumococcal polysaccharides were assessed on weeks 0, 2 and 4 post vaccination. Sera collected 4-weeks post vaccination were used to assess (B) the binding efficacy of antibodies to *S. pneumoniae* using flow cytometry and (C) the ability of antibodies to induce opsonophagocytic killing by PMNs. (A) Data are presented as percentage of standard sera and are pooled from a total of 3 experiments with a total of n=9 mice per group per timepoint. * denotes significant differences between PMNs– & Tregs-depleted mice and group matching the symbol color as determined by Kruskal-Wallis test followed by Dunn’s multiple comparisons test. (B) Data are from 5 separate experiments with each dot representing the average of technical replicates. * denotes significant differences between the indicated groups as determined by One-way ANOVA followed by Šídák’s multiple comparisons test and # denotes significant differences from naïve as determined by One-way ANOVA followed by Holm-Šídák’s multiple comparisons test. *** is *p*≤0.001, and # is *p*≤0.001. Bar graphs represent the mean +/− SD. (C) Data are pooled from 3 separate experiments with each dot representing the average of technical replicates from an individual mouse. * denotes significant differences between the indicated groups as determined by One-way ANOVA followed by Šídák’s multiple comparisons test. * is *p*≤0.05 and ** is *p*≤0.01. All line graphs represent the mean +/− 95% confidence interval and all bar graphs represent the mean +/− SD.

To confirm these findings, we used a different model of Treg depletion for assessment of antibody responses to PCV using diphtheria toxin (DT) mediated depletion (Jin et al., 2017). For that, we used DT injections to deplete Tregs either alone or in conjunction with PMN depletion in DEREG mice (Lahl et al., 2007) (Fig. S21A). We achieved ∼25% depletion of Tregs in the spleen (Fig. S21B). Treg depletion in PMN-depleted DEREG mice increased their IgG levels 4 weeks post vaccination compared to vehicle treated PMN-depleted mice (Fig. S21C). The above data suggest that the increase in Tregs seen in PMN-depleted mice contributes to the defective antibody response observed in these mice as reduction of the Tregs rescues the levels and function of the antibodies to similar levels seen in PMN-sufficient vaccinated mice and reduction of Tregs alone in PMN-sufficient vaccinated mice has no effect.

### Reducing Tregs in PMN-depleted hosts rescues the protective capacity of the sera against pneumococcal infection

Finally, we assessed the protectiveness of the sera from PMN– and/or Treg-depleted mice *in vivo* against infection by adoptively transferring sera from each group into naïve mice. Recipient mice were then infected 1hr later with 3×10^5^ CFU of *S. pneumoniae* TIGR4 strain, assessed for blood bacterial burden 48 hours post infection, and monitored for survival. Recipients of sera from vaccinated PMN-sufficient mice were able to control bacterial numbers better than the ones receiving sera from vaccinated PMN-depleted mice with 50% and 87.5% of the mice becoming bacteremic respectively (Fig. 5A). Receiving sera from PMN– & Treg-depleted mice rescued the ability of the recipients to control infection with only 25% of the mice becoming bacteremic (Fig. 5A). Additionally, receiving sera from Treg-depleted mice did not affect control of blood bacterial burden (Fig. 5A). Looking at survival, only 10% of recipients of sera from naïve mice survived the infection compared to ∼70% of the recipients of sera from vaccinated PMN-sufficient mice (Fig. 5B). Receiving sera from PMN-depleted mice resulted in only 30% survival, compared to ∼81% survival of mice that received sera from PMN– & Treg-depleted mice (Fig. 5B). The above data suggest that reduction of Tregs in PMN-depleted mice rescues the protectiveness of the sera to levels similar to PMN-sufficient vaccinated mice.

**Figure 5:**
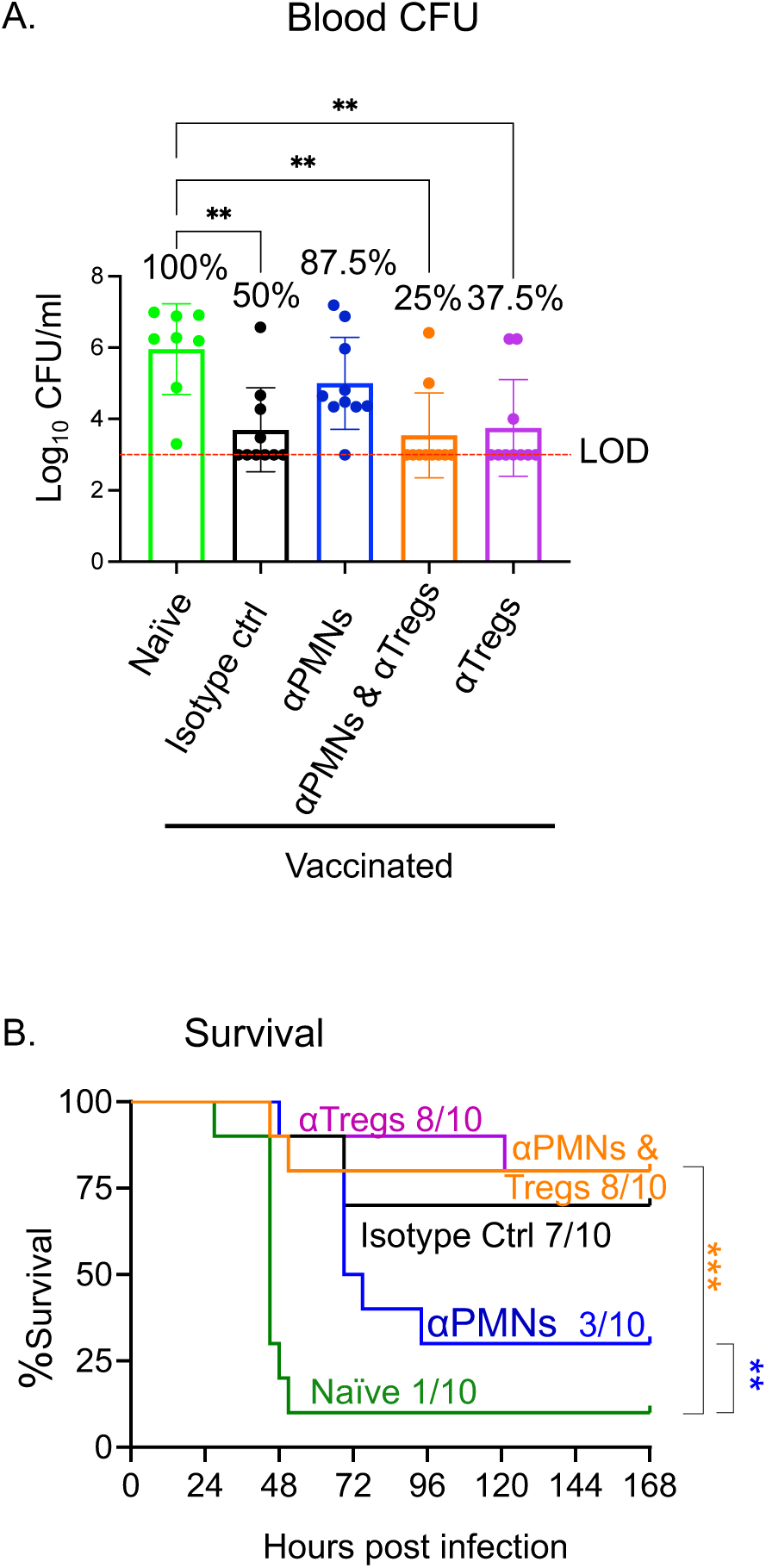
Effect of Treg depletion on the protective efficacy of sera against pneumococcal infection. Adult C57BL/6 female mice were injected with PCV or PBS and depleted of their PMNs and/or Tregs (αCD25) or treated with an isotype control. Sera were collected from all mice 4 weeks post vaccination. Sera from each mouse group was adoptively transferred into a new set of naïve mice which were subsequently infected with 3×10^5^ CFU of *S. pneumoniae*. (A) Bacterial burden in the blood was assessed 45 hours post infection. (B) All mice were monitored for survival for 7 days following infection. (A) Data are pooled from 2 separate experiments and each dot represents an individual mouse with a total of n=10 mice per group. * denotes significant differences between the indicated groups as determined by Kruskal-Wallis test followed by Dunn’s multiple comparisons test. Bar graphs represent the mean +/− SD. (B) Data are pooled from 2 separate experiments with a total of n=10 mice per group. * denotes significant differences between the indicated groups as determined by Mantel-Cox test. ** is *p*≤0.01, and ** is *p*≤0.01.

### Vaccination of human participants alters the phenotype of circulating PMNs

As PMN responses vary significantly across species (Nauseef, 2023), we wanted to test the clinical relevance of these findings. To do so we recruited unvaccinated adult female participants, administered PCV to them, and monitored the phenotype of PMNs in the circulation over time in response to vaccination by examining the expression of T cell engaging and inhibiting markers using flow cytometry (Fig. S22). To test if PMNs respond to PCV and alter their phenotype, we used the OMIQ software to perform opt-SNE analysis to visualize the data, and cluster them using FLOWSOM. We found that PMNs separated into different clusters with varying levels of maturity (based on CD10 expression, Fig. 6A and S23) with cluster 6 being highest followed by 3, 2, 4, 1, 9, 7, 5, and 8 the lowest. Clusters 6 (most mature) was the only cluster that expressed PDL-1 and had high levels of all the other markers, while cluster 8 (least mature), did not express any of the markers we tested (Fig. 6A, 6C and S23). Upon vaccination, we observed a shift in the abundance of circulating clusters where cluster 1, 2 and 6 decreased in abundance, cluster 3, 4 and 5 increased, while cluster 7 and 8 did not change (Fig. 6B). The clusters that decreased from the circulation expressed the highest levels of CD39, ARG1 and HLADR (Fig. 6C) and may be homing to the tissues as we observed in mice. When we examined the expression levels of the various markers at week 1 following vaccination, we found that expression of HLADR, CD86, CD39 and PDL-1 increased on all the clusters that expressed them, CD73 either increased or decreased depending on the cluster, while ARG1 decreased (Fig. 6C). Overall, these findings suggest that in response to PCV in humans, the phenotype of circulating PMNs changes, where particular clusters increase expression of T cell engaging and suppressing markers and possibly exit from the circulation into tissues.

**Figure 6:**
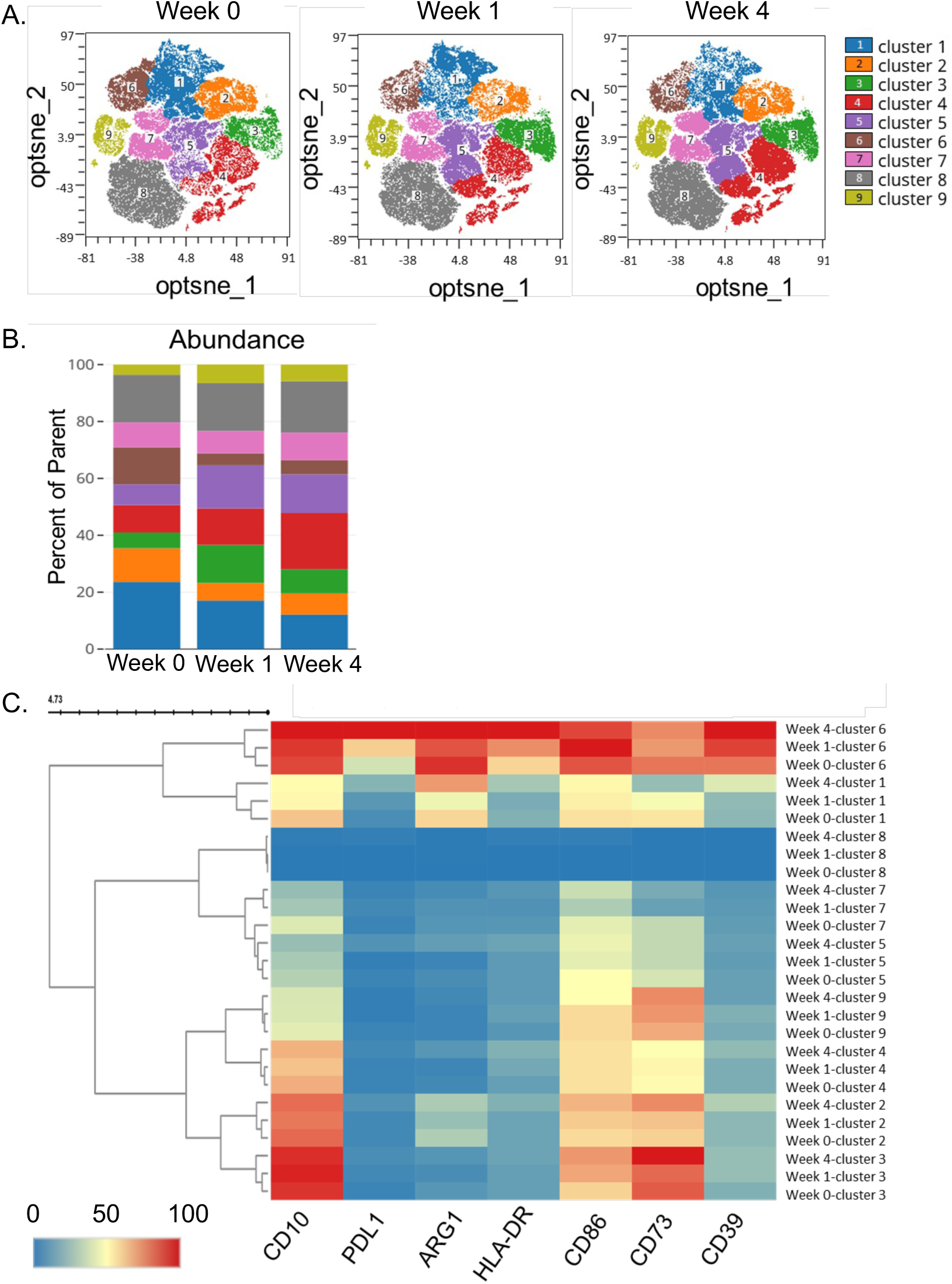
PMN phenotype following PCV vaccination in humans. Young (24-29 years old) female donors were vaccinated with PCV. Blood was collected prior (week 0) to and on weeks 1 & 4 post vaccination. PMNs were isolated from peripheral blood and assessed using flow cytometry. Expressions of HLA-DR, CD10, CD86, PD-L1, ARG1, CD73 and CD39 on PMNs were determined. (A) Clustering of investigated markers throughout the weeks, (B) cluster abundance throughout the weeks, and (C) heat map of markers throughout the different weeks and clusters were analyzed using FLOSOM via OMIQ. Data are pooled from six donors.

### PMNs from PCV immunized donors inhibit FOXP3^+^ T cell expansion *in vitro*

Finally, we tested whether PMNs from vaccinated participants can suppress FOXP3^+^ CD4^+^ T cells. For that, at one week post vaccination, a timepoint where we observed phenotypic changes in PMNs in response to vaccination (Fig. 6), we performed an *in vitro* assay where PBMCs from each participant were cocultured with or without PMNs (from the same donor) and stimulated with PCV for 48 hours and T cells assessed (Fig. S24). Similar to what we found in the preclinical model, the presence of PMNs resulted in reduction of FOXP3^+^ T cells. Addition of PMNs to PBMC cultures lead to a decrease in the percentages of total and antigen experienced FOXP3^+^ T cells in 67% of participants (Fig. 7A). We also observed a decrease in Ki67^+^ antigen experienced FOXP3^+^ T cells in the presence of PMNs in 100% of the participants (Fig. 7B), suggesting that PMNs were suppressing proliferation. These data suggest that PMNs inhibit FOXP3^+^ T cell responses in humans following PCV administration.

**Figure 7:**
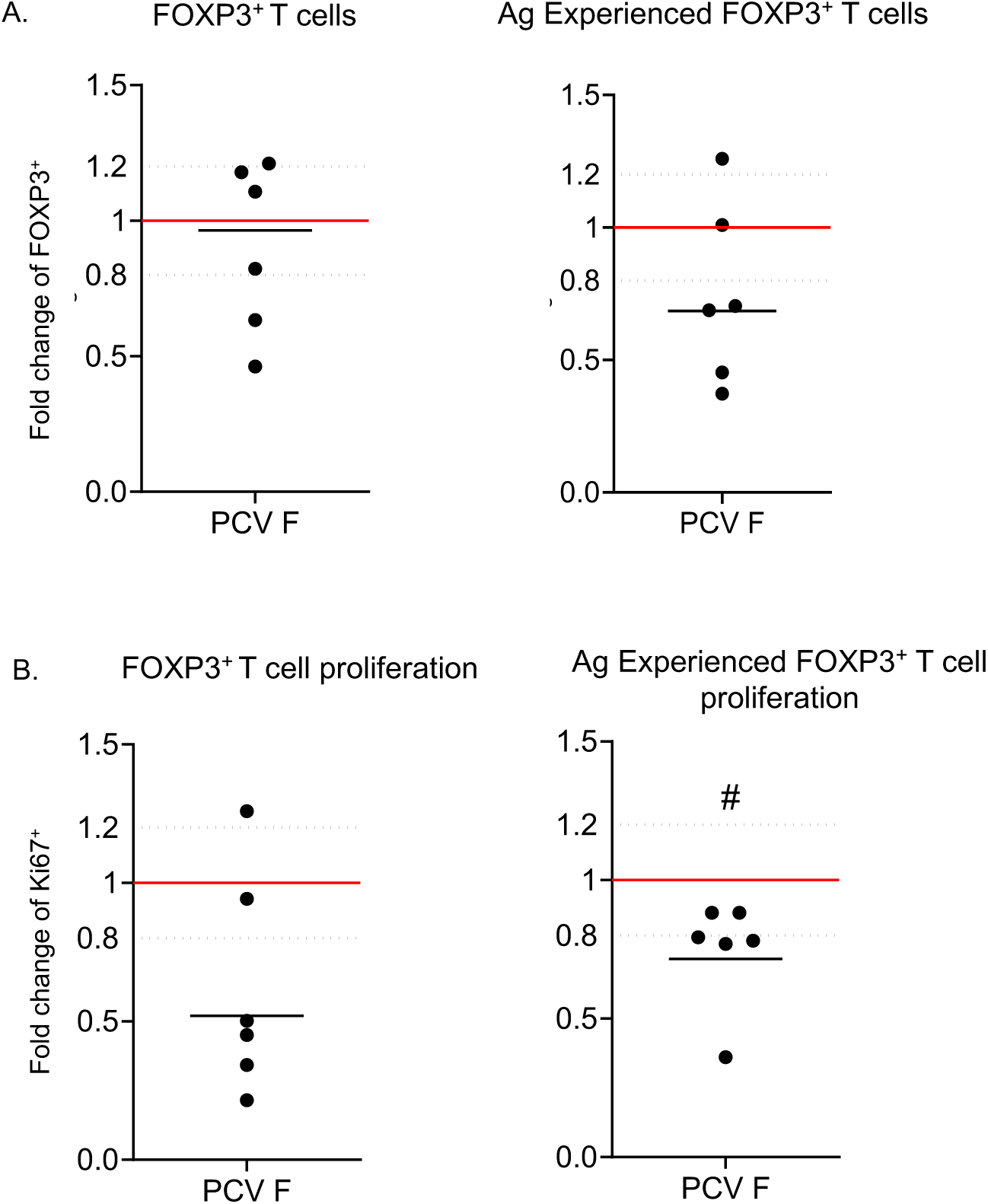
Human PMN and PBMC co-cultures following PCV vaccination. Young (24-29 years old) female donors were vaccinated with PCV. One week post vaccination, PBMCs and PMNs were isolated from peripheral blood. PBMCs were cultured in the presence or absence of matched donor PMNs for 2 days with PCV as a stimulant. Fold change of the percentages of (A) FOXP3^+^ and (B) Ki67^+^ FOXP3^+^ cells gated from total (left) and antigen experienced (right) CD4^+^ T cells are shown. Data are pooled from six donors and each dot represents a donor. Data are presented as fold change of PBMC divided by PBMC +PMNs conditions. The dashed lines represent a 20% difference in responses between the conditions. Graphs represent the mean, and # indicate significant differences from 1 as measured by a one-sample t-test.

## DISCUSSION

There are a growing number of studies investigating the effect of neutrophils on adaptive immune responses with a few of them focusing on vaccine responses (Bhattacharya et al., 2020; Musich et al., 2018; Trentini et al., 2016). In this study, we report a new mechanism by which PMNs control antibody responses to the clinically relevant PCV by controlling Treg numbers. In fact, when neutrophils were depleted at the time of PCV vaccination, there was an expansion in the number of Tregs that resulted in a defect in protective antibody responses (Tchalla et al., 2020). Reduction of Tregs in PMN-deficient vaccinated mice boosted antibody number, function, and protective efficacy in the sera. This work demonstrates that PMNs drive production of functional antibodies against pneumococci in response to PCV by limiting the expansion of immune suppressive Tregs.

While the effect of PMNs on Tregs has not been investigated in the context of clinically relevant vaccines, several studies have looked at the relationship between Tregs and PMNs in models of inflammation and disease (Gao et al., 2015; Himmel et al., 2011; Lewkowicz et al., 2013; Mishalian et al., 2014; Nadkarni et al., 2016; Perobelli et al., 2016; Richards et al., 2010). Here we report that PMNs can inhibit Tregs, which is in contrast to prior work demonstrating that PMNs promote Tregs in models where high level of inflammation is induced (Gao et al., 2015; Mishalian et al., 2014; Perobelli et al., 2016). The exact way by which PMNs inhibit Tregs in response to PCV remains to be elucidated, however multiple mechanisms could be involved. Some possibilities could be via control of the transcription factor FOXP3, a master regulator of Treg induction and function (Fontenot et al., 2003; Ono, 2020) or control of cytokines required for Treg differentiation and viability. In traumatic brain injury patients, release of serine proteases (neutrophil elastase and proteinase 3) by infiltrating PMNs cleaved cytokine receptors on the surface of T cells including IL-2Rα, the IL-2 receptor subunit that is important for IL-2 signaling in Tregs (Bank et al., 1999; Harris et al., 2023). In addition, those proteases can cleave cytokines themselves including TGFβ and IL-2, leading to limited activation of T cell response (Ariel et al., 1998; Fu et al., 2020). In response to PCV, PMNs may similarly inhibit Treg induction by release of their proteases to deplete Treg-inducing cytokines and/or their respective receptors on the surface of Tregs. Signaling of retinoic acid (RA) in naïve T cells enhances TGFβ-mediated induction of Tregs (Ohkura et al., 2011) and RA boosts PMN anti-bacterial functions (Stream et al., 2024). It is possible that RA sequestration by PMNs limits its availability for Treg induction. PMNs could also be inhibiting Tregs through release of their granule component cathelicidin antimicrobial peptide, the human form of which (LL-37) was shown to induce Treg apoptosis *in vitro* (Mader et al., 2011). Alternatively, PMNs could be suppressing Tregs through the release of miRNA containing extracellular vesicles (EVs). PMN-derived EVs containing miRNAs have been shown to modulate responses in other cells including macrophages and endothelial cells (Glemain et al., 2022; Gomez et al., 2020; Jiao et al., 2021). The absence of miRNA-146a has been shown to promote Treg expansion, and other miRNAs are known to control Treg induction (Kunze-Schumacher and Krueger, 2020; Lu et al., 2021a). PMNs, which express miRNA-146a among others (Arroyo et al., 2021; Garley et al., 2024), could use such mechanism to limit Treg expansion. All these potential mechanisms can be avenues of future research for this work.

The findings here suggest that the expanded Tregs in PMN-depleted mice are detrimental and contribute to the defective antibody responses. In general, Tregs have been shown to negatively affect immune responses to vaccines. Depletion of Tregs using αCD25 in vaccination models against herpes simplex virus (HSV) and *Plasmodium berghei*, increased CD8^+^ T cell responses to their respective vaccine epitopes and led to increase protection against subsequent challenge with the respective pathogens (Moore et al., 2005; Toka et al., 2004). In humans, responses to an experimental vaccine of dendritic cells loaded with HIV peptides (Levy et al., 2014) found that patients with lower levels of HIV-specific Tregs displayed better T-effector cell responses following vaccination (Brezar et al., 2015). In the context of *S. pneumoniae*, blocking of CTLA-4, an effector molecule on Tregs, enhanced levels of antibodies against pneumococcal polysaccharides in immunized mice and higher Tregs in older adults were associated with decreased proliferation of CD4^+^ effector T cells in response to the pneumococcal protein AliB (Boudewijns et al., 2005; He et al., 2021). Additionally, Tfrs, which we found are increased in PCV-vaccinated PMN-depleted mice, have been shown to inhibit B cells responses and promote the contraction of GC reactions (Jacobsen et al., 2021; Sage et al., 2016). The findings here, showing an improvement of vaccine response upon depletion of the expanded Tregs in PMN-depleted mice, are in line with the reported inhibitory role of Tregs in vaccine responses.

There have been several studies examining how Tregs and Tfrs inhibit B cell responses (Wang and Zheng, 2013). They can affect B cell number, differentiation, and antibody responses via release of soluble mediators such as the neuropeptide neuritin, IL-10 and TGFβ or through contact dependent mechanisms involving CTLA4, PD1 and granzymes (Gonzalez-Figueroa et al., 2021; Tan et al., 2022; Wang and Zheng, 2013). This suppression can lead to a decrease in B cell expression of activation-induced deaminase (AID) (Lim et al., 2005). However, B cell numbers and AID expression were not affected in this study. It is possible that the DNA binding capacity or the enzymatic activity of AID are affected instead. AID activity is regulated by its phosphorylation status which can either enhance or inhibit its enzymatic activity, depending on the amino acid being phosphorylated (McBride et al., 2008; Park, 2012) by protein Kinase A (PKA) (Basu et al., 2005; Chaudhuri et al., 2004; Demorest et al., 2011; Gazumyan et al., 2011). It is therefore possible for Tregs to be regulating AID phosphorylation in response to PCV, rather than overall expression through modulation of cAMP as they have been shown to do in DCs and effector T cells (Klein and Bopp, 2016).

We found that the phenotypes observed in the preclinical model are also found in human participants, highlighting the clinical relevance of this work. Upon vaccination of healthy human donors, PMNs in the circulation changed phenotype suggesting they are responsive to vaccination. Using peripheral blood collected from donors after PCV vaccination for *in vitro* cocultures, addition of PMNs to PBMCs lead to a reduction in FOXP3^+^ T cells and their proliferation, recapitulating the Treg inhibiting property of PMNs seen in the mouse models. However, further clinical studies are needed to assess the *in vivo* role of PMNs in protecting human hosts against pneumococcal disease, including linking PMN phenotype to antibody responses. Nevertheless, these findings can impact the design of improved pneumococcal vaccine formulations that offer better efficacy, especially in the more vulnerable populations. Despite their availability, pneumococcal vaccines do not mitigate the growing global burden of *S. pneumonia* infections (CDC, 2022; Wiemken et al., 2014; Wiese et al., 2016). This is due in part to the fact that the vaccines do not protect all populations equally, where male donors and aged individuals mount lower responses compared to female counterparts. Interestingly, Tregs are reported to be higher in males compared to sex-matched females and higher in older adults compared to younger people (Afshan et al., 2012; Raynor et al., 2012; Robinson et al., 2022). In this work, we identified the role of the PMN-Treg axis in controlling antibody responses in females that can be targeted for improvement of current vaccine formulations moving forward. While further studies are needed to tease apart the specific mechanisms behind this inhibition of Tregs by PMNs, the findings have important implications for development of future vaccine strategies to improve efficacy to infections. Indeed, this PMN-Treg axis can be further studied in the context of sex and age-driven differences in vaccine responses (Flanagan et al., 2017; Pollard and Bijker, 2021) and can be targeted to increase vaccination efficacy in the vulnerable older adults and male populations. The exact mechanisms of Treg inhibition by PMNs need to be elucidated to pinpoint what pathways or molecules to target specifically in the future.

### Limitations of the Study

One limitation of the study is the focus on young adult females, both in preclinical mouse work and in human participants. Neutrophil responses are known to be affected by both host age and biological sex (Klein and Flanagan, 2016; Lu et al., 2021b; Simmons et al., 2021; Spitzer, 1999). Therefore, their role in vaccine-mediated responses may be different in the more vulnerable male and aged hosts. Indeed, our prior work in preclinical models (Tchalla et al., 2024) as well as data from human studies (Soneji and Metlay, 2011; Wagenvoort et al., 2017; Wiese et al., 2016) found that PCV efficacy is lower in males compared to females and declines with host age. Therefore, we decided to first focus on the hosts that are the most protected following vaccination (young adult females) and characterize PMN responses in them first and move to less protected hosts as a next step. Currently ongoing studies in our lab are exploring the effect of aging and biological sex on PMN responses to PCV. Another limitation is, for human participants, we are limited by access to cells in the circulation. Therefore, and are unable to examine the effect of PMNs on secondary lymphoid organs. We attempted to address that by performing PMN/PBMC co-cultures and were able to recapitulate the inhibitory effect of PMNs on Tregs that we observed *in vivo* in preclinical studies.

Further by being limited to responses in the circulation, we are unable to test if PMN phenotype is altered upon tissue entry in people. However, unlike in mice, the phenotype of circulating PMNs in human participants did change in response to PCV administration, emphasizing differences in PMN behaviors across species. Finally, although we identified that control of Tregs by PMNs is one of the mechanisms by which they affect antibody responses to PCV, how they do so remain an open question. In summary, despite the discussed limitations, this study revealed new insights into how innate immunity controls adaptive responses, which has implications for future vaccine design, highlighting the importance of taking PMN responses into consideration.

## RESOURCE AVAILABILITY

### Lead contact

Requests for further information and resources should be directed to and will be fulfilled by the lead contact, Elsa Bou Ghanem.

### Materials availability

The study did not generate new unique reagents.

### Data and code availability

All data reported in this paper will be shared by the lead contact upon request. This paper does not report original code. Any additional information required to reanalyze the data reported in this paper is available from the lead contact upon request.

## ACKNOWLEDGMENTS

This work was supported by National Institute of Health grant R01 AG068568-01A1 to ENBG, Mark Diamond Research Funds to EYIT, and an AHA predoctoral fellowship to AB. The content is solely the responsibility of the authors and does not necessarily represent the official views of the National Institutes of Health.

## AUTHOR CONTRIBUTIONS

EYIT conducted research, acquired data, analyzed data, and wrote the manuscript. AB conducted research, acquired data, and analyzed data. AYK, EL, MB, MC conducted research and acquired data. EAW provided materials. ENBG designed the research studies, wrote, and edited the manuscript, and had responsibility for final content. All authors read and approved the manuscript.

## SUPPLEMENTAL INFORMATION

Document S1. Figures S1–S24.

## MATERIALS AND METHODS

### Experimental Model and Study Participant Detail

#### Mice

Adult (10-12 weeks old) female C57BL/6 mice were purchased from The Jackson Laboratory (Bar Harbor, Maine). Our study focused on female hosts, as there is a well-documented sex-based difference in responses to PCV where females elicit higher antibody responses and are better protected from subsequent infection compared to males (de St Maurice et al., 2016; Tchalla et al., 2024; Wagenvoort et al., 2017). Therefore, we examined the mechanisms behind vaccine efficacy in the better protected hosts. Ongoing work in our lab is focused on sex-based difference in PMN responses in vaccinated hosts (Tchalla et al., 2024) which is beyond the scope of this study. Adult (10-16 weeks old) C57BL/6-Tg(Foxp3-DTR/EGFP)23.2Spar/Mmjax DEREG mice were a kind gift from Elizabeth Wohlfert and have been previously described in detail (Jin et al., 2017; Lahl et al., 2007). All mice were housed in a specific-pathogen-free environment at the University at Buffalo Jacobs School of Medicine and Biomedical Sciences animal facility.

### Human participants

Healthy unvaccinated female participants (24-29 years old) were recruited. Individuals who were pregnant, had acute infections within the last 2 weeks, took medication 48 hours prior, had history of chronic infections, had conditions that altered immunity (autoimmunity, on immune modulating drugs, history of cancer) were excluded from the study. At the first visit (week 0), donors had their blood drawn then were administered the Polysaccharide Conjugate Vaccine Prevnar 20 (Pfizer, Wyeth Pharmaceuticals) intramuscularly as per manufacturer’s instructions. At week 1 and 4 following vaccination, donors had their blood drawn for subsequent PMN phenotyping. Blood from week 1 was also used for coculture assays. All blood was collected into tubes containing sodium citrate and visits occurred in the morning (8AM-10AM) to control for circadian variations in PMN phenotype (Casanova-Acebes et al., 2013). All enrolled participants signed approved informed consent forms.

### Study approvals

All work with mice was performed in accordance with the recommendations in the Guide for the Care and Use of Laboratory Animals published by the National Institutes of Health. All procedures were reviewed and approved by the University at Buffalo Institutional Animal Care and Use Committee (IACUC), approval number MIC33018Y. All work with human donors was approved by the University at Buffalo Human investigation Review Board (IRB), approval number STUDY00007111.

### Method Details

#### Murine immunization and sera collection

Mice were vaccinated intramuscularly with 50ul of the Polysaccharide Conjugate Vaccine Prevnar 13 (Wyeth Pharmaceuticals) into the caudal thigh muscle. 10ul of blood from each mouse was collected from the tail at weeks 0, 2 and 4 following vaccination and used for assessment of antibody titers. Sera from each mouse group were collected via cardiac puncture 4 weeks following vaccination, pooled and aliquoted to assess antibody quantity and functions in subsequent assays.

### Bacteria

*Streptococcus pneumoniae* TIGR4 strain (serotype 4) was a kind gift from Andrew Camilli. Bacteria were grown to mid-exponential phase in Todd–Hewitt broth (BD Biosciences) supplemented with Oxyrase at 37°C/5% carbon dioxide, aliquoted in growth media supplemented with 20% glycerol and saved at –80°C as previously described (Tchalla et al., 2024; Tchalla et al., 2020). Prior to use, bacteria were thawed on ice, washed, and enumerated by serial dilution in PBS followed by dribbled plating on Tryptic Soy Agar (TSA) plates supplemented with 5% sheep blood (Northeast Laboratory Services).

### Murine neutrophil depletion

Mice were injected intraperitoneally with 50μg of the Ly6G-depleting antibody IA8 or isotype IgG control following the same schedule as shown in Fig. S2A and previously described (Tchalla et al., 2020). PMN depletion in the spleen was assessed by a myeloperoxidase ELISA (Invitrogen) as per manufacture’s instruction (Tchalla et al., 2020).

### Murine regulatory T cell depletion

B6 mice were treated intraperitoneally with 300μg of the CD25-depleting antibody PC-61.5.3 or isotype IgG1 control following the schedule shown in Fig. S19A. DEREG mice were treated intraperitoneally with 1μg of diphtheria toxin (Millipore) or DMSO as vehicle control following the schedule shown in Fig. S21A. Depletion efficiency was assessed by flow cytometry (Fig. S19 & S21B).

### Murine organ/tissue processing and flow cytometry

Blood, spleens and/or vaccine draining lymph nodes (vLNs-inguinal and popliteal) were harvested from euthanized mice. Single cell suspensions from spleens and vLNs were prepared in RPMI 1640 media supplemented with 10% Fetal Bovine Serum. Following red blood cells lysis with an in-house hypotonic solution (8.29g ammonium chloride, 1g sodium bicarbonate, 0.038g trypsin/ethylenediaminetetraacetic acid (EDTA) and 1L of water), the remaining cells were enumerated, incubated with Fc block (2.4G2), Live/Dead dye (Invitrogen) and stained with additional antibodies in separate panels.

For PMNs, cells were stained with Ly6G (1A8), CD11b (M1/70), CXCR4 (2B11), CCR7 (4B12), CD80 (16-10A1), CD86 (GL1), CD11c (N418), MHC-II (M5/114.15.2), PD-L1 (MIH5), CD39 (24DMS1) and/or CD73 (TY/11.8) antibodies followed by permeabilization with Cytofix/Cytoperm (BD) and intracellular staining with ARG1 (A1exF5).

For B cells, cells were stained with CD45R/B220 (RA3-6B2), CD138 (281-2), CD95 (SA367H8), and CD38 (90) antibodies followed by permeabilization with Fix/Perm buffer from the FOXP3/Transcription factor staining buffer set (eBioscience) and intracellular staining with AID (Polyclonal, Bioss).

For T cells, cells were stained with CD4 (RM4-5), TCRβ (H57-597), CD11a (M17/4), CD49d (R1-2), PD1 (J43) and/or CXCR5 (SPRCL5) antibodies followed by permeabilization with Fix/Perm buffer from the FOXP3/Transcription factor staining buffer set (eBioscience) and intracellular staining with FOXP3 (FJK-16s), RORγT (Q31-378), Tbet (eBio4B10) and GATA3 (L580-823).

For intracellular cytokine staining in PMNs, cells were incubated in Brefeldin A in RPMI 1640 media supplemented with 10% Fetal Bovine Serum and 1% Pen/Strep and stimulated with 9×10^6^ CFU/well of heat-killed *S. pneumoniae* TIGR4 strain, 10μl/well of PCV, 100nM of phorbol-12-myristate 13-acetate (PMA) or media alone for 3 hours. Cells were then washed and stained extracellularly with Fc block, Live/Dead dye, Ly6G, and CD11b antibodies followed by intracellular staining with IFNγ (XMG1.2) after permeabilization with Cytofix/Cytoperm (BD). For intracellular cytokine staining in T cells, cells were incubated in Brefeldin A (BD) in RPMI 1640 media supplemented with 10% Fetal Bovine Serum and 1% Pen/Strep and stimulated with a mix of 81nM of PMA and 1.34μM of Ionomycin (Cell activation cocktail, Biolegend) or media alone for 2 hours. Cells were then washed and stained extracellularly with Fc block, Live/Dead dye, CD4, TCRβ, CD44 (IM7) and LAP (TW7-16B4) antibodies followed by intracellular staining with IFNγ (XMG1.2), IL-17A (TC11-18H10), IL-4 (11B11) and IL-10 (JES5-16E3) antibodies after permeabilization with Fix/Perm buffer from the FOXP3/Transcription factor staining buffer set (eBioscience). Following staining, cells were washed with FACS buffer (HBSS, 1% FBS and 0.1% sodium azide) and fluorescence was assessed using a BD Fortessa. At least 20,000 cells in the live, CD4^+^TCRβ^+^ and B220^+^ cells gates respectively were collected and analyzed using the FlowJo software.

### Murine antibody enzyme-linked immunosorbent assay

Antibodies in sera were assessed using an ELISA assay as previously described (Bhalla et al., 2021; Tchalla et al., 2020). Briefly, Nunc-maxisorp plates (Thermo Scientific) were coated at 4°C overnight with type 4 Pneumococcal Polysaccharide (ATCC®) at 2μg/well. The next day, plates were washed in PBS supplemented with 0.25% Tween (PBS/T) and sera preabsorbed with a pneumococcal cell wall polysaccharide mixture (CWP-multi, Cederlane) were added to the plates. After a 2-hour incubation, plates were washed and incubated with diluted horseradish peroxidase (HRP) conjugated anti-mouse IgM (Invitrogen), IgG (Millipore Sigma) or IgG1, IgG2b, IgG2c or IgG3 (Southern Biotech). One hour later, plates were washed, TMB substrate (Thermo Scientific) was added, and absorbance was read at 650nm using a Biotek microplate reader for 10 minutes at 1-minute intervals. Antibody units were calculated as a percentage of standard sera included in each assay. Standard sera were obtained from mice that were intranasally inoculated with *S. pneumoniae* serotype 4 weekly for 3 weeks and immunized with PCV at week 4.

### Murine IFNγ enzyme-linked immunosorbent assay

Single cell suspension of splenocytes were prepared in 2ml of RPMI 1640 media supplemented with 10% Fetal Bovine Serum. IFNγ concentrations were quantified using the Mouse IFNγ ELISA kit (Invitrogen, KMC4021) per manufacturer’s instructions.

### Murine IFNγ blocking *in vivo*

Mice were treated intraperitoneally with 200μg of the IFNγ-blocking antibody XMG1.2 or isotype control following same schedule as neutrophil depletion shown in Fig. S2A.

### Murine cytokine/chemokine array

Pan cytokines were assessed using the Proteome Profiler Mouse XL Cytokine Array kit (R&D, ARY028) following the manufacturer’s instructions as previously described (Tchalla et al., 2024). Briefly, single cell suspension of splenocytes were prepared in 2ml of RPMI 1640 media supplemented with 10% Fetal Bovine Serum. Cell suspensions were aliquoted and saved at – 80°C. Samples were thawed on ice prior to use, lysed with Triton-X (VWR, 0694) and quantified for protein concentrations using the Pierce BCA protein assay kit (Thermofisher, A55864) per the manufacturer’s instructions. All samples were brought to the same protein concentrations and used for pan-cytokine assessment. Protein membranes were read using the BioRad ChemiDoc MP Imaging System and the intensity for each dot on the blots was quantified using the QuickSpots Image Analysis Software. The intensity values of each blot were blanked using their respective buffer only controls. Intensity values across multiple blots were normalized using the blot with the highest standard. Fold change of the values from vaccinated PMN-depleted mice over those of vaccinated PMN-sufficient mice were calculated and log transformed.

### Murine splenocyte culture assays

For mice, single cell suspension of splenocytes were prepared in 2ml of RPMI 1640 media supplemented with 10% Fetal Bovine Serum one day post vaccination. To remove PMNs from the obtained suspensions, splenocytes were stained with a PE labeled anti-Ly6G antibody and treated using the EasySep™ Release Mouse PE Positive Selection Kit (StemCell) following manufacturer’s instruction. One million cells per conditions were used and treated with 5×10^7^ CFU heat-killed *S. pneumoniae* or 1μl/ml αCD3 (positive control). Cells were cocultured for two days before assessing percentages of Tregs.

Antibody binding to the surface of *S. pneumoniae*

Sera collected 4-weeks post vaccination were used to assess antibody binding to *S. pneumoniae* (TIGR4) as previously described (Tchalla et al., 2024; Tchalla et al., 2020). After 30 minutes of incubation at 37°C, cells were washed with FACS buffer and labeled with an APC-tagged anti-mouse IgG (polyclonal) antibody for 30 minutes on ice in the dark. Cells were then washed, resuspended in FACS buffer and fluorescence was assessed using a BD Fortessa. At least 10,000 cells were analyzed using the FlowJo software.

### Murine opsonophagocytic killing assay (OPH)

Neutrophils were isolated from the bone marrow of mice using density centrifugation with Histopaque 1119 and Histopaque 1077 (Sigma) as previously described (Bhalla et al., 2021; Bou Ghanem et al., 2015; Tchalla et al., 2020). Briefly, Neutrophils were resuspended in HBSS (without Ca^2+^ or Mg^2+^) media supplemented with 0.1% gelatin and used in the assay. Sera collected 4-weeks post vaccination were incubated with *S. pneumoniae* TIGR4 for 5 minutes and added to neutrophils in HBSS (with Ca^2+^ and Mg^2+^). Following a 40-minute incubation at 37°C, reactions were plated on TSA plates with 5% sheep blood. Percent killing was calculated relative to a no neutrophils control for each sera condition.

### Murine adoptive transfer of sera

Four weeks following immunization, naïve, vaccinated neutrophil– and Treg-sufficient, neutrophil-depleted, neutrophils– and Treg-depleted and Treg-depleted were euthanized. Blood was harvested via cardiac puncture; sera were obtained from the blood and pooled for each mouse group. A volume of 250μL of sera was transferred intraperitoneally into naive recipients which were infected 1hour later.

### Animal infections

Mice were infected intratracheally with 3×10^5^ CFU of *S. pneumoniae* serotype 4 (TIGR4) using the tongue pull method as previously described (Lenhard et al., 2022; Tchalla et al., 2020). Mice were monitored for survival for a total of 7 days post infection and assessed daily for bacteremia by plating blood collected in 50mM EDTA on blood agar plates for enumeration of bacterial CFU.

### Human PMN isolation and flow cytometry

For participants, PMNs were isolated from peripheral blood of week 0, 1, 4 post vaccinated donors using EasySep™ Direct Human Neutrophil Isolation Kit (StemCell) respectively following manufacturers’ instructions. For each donor and visit 1×10^5^ PMNs were added to each well and then stained with CD16 (eBioCB16), CD14 (61D3), CD10 (MEM-78), HLA-DR (L243), CD86 (BU63), PD-L1 (MIH1), CD73 (AD2), and/or CD39 (eBioA1) antibodies followed by permeabilization with Cytofix/Cytoperm (BD) and intracellular staining with ARG1 (A1exF5). Following staining, cells were washed with FACS buffer and fluorescence was assessed using a BD Fortessa. At least 5000 cells in the CD16^+^ gate were collected and analyzed using the FlowJo software.

### Human PMN FLOWSOM analysis

Analysis and data visualization of the PMN phenotype with and without vaccination over time was completed using a software OMIQ (Dotmatics). Traditional gating was done to differentiate the neutrophil population and all the markers. Dimensional reduction was next performed with optimized parameters for T-distributed stochastic neighbor embedding which improves visualization of large datasets by reducing it to 2D/3D map while preserving local structure. Clustering into neutrophil subsets was done using Flow-cytometry specific self-organizing map (FLOWSOM)(Van Gassen et al., 2015) algorithm. Clustered heat maps were made based on meta-cluster gating to characterize the different PMN subsets based on the expression of different markers.

### Human PMN/PBMC coculture assays

For participants, PBMCs and PMNs were isolated from peripheral blood of 1-week vaccinated donors using the One-Step Polymorph (Accurate Chemical) and EasySep™ Direct Human Neutrophil Isolation Kit (StemCell) respectively following manufacturers’ instructions. For each donor, 2×10^5^ PBMCs were incubated in (RPMI with 10% FBS) with or without addition of 5×10^4^ PMNs and stimulated with 10μl of PCV. Cells were cocultured for two days before assessing percentages and proliferation of T cells. At 48 hours post incubation, T cells were stained with CD11a (HI111), PD1(EH12.1), CD4(S3.5), CD49d (9F10) and CXCR5 (MU5UBEE) followed by permeabilization with Fix/Perm buffer from the FOXP3/Transcription factor staining buffer set (eBioscience) and intracellular staining with FOXP3 (PCH101), RORγT (AFJKS-9), Tbet (eB4B10), GATA3 (L580-823) and Ki67 (B56). Following staining, cells were washed with FACS buffer and data collected using a BD Fortessa. At least 10,000 cells in the live CD4^+^ gate were collected and analyzed using the FlowJo software.

### Quantification and statistical analysis

All statistical analysis was done using GraphPad Prism version 11 software. Data were checked for normality distribution using the Shapiro-Wilk test. Significant differences were determined by One-way analysis of variance followed Šídák’s, Holm-Šídák’s, Tukey’s or Dunnett’s multiple comparisons test; Kruskal-Wallis One-way analysis of variance followed Dunn’s multiple comparisons; Student’s t test; or one sample t-test as appropriate. Survival was analyzed using the log-rank (Mantel-Cox) test. All *p* values <0.05 were considered significant.

