## Supplemental Data 1 for "Neutrophils induce effective antibody responses to the pneumococcal conjugate vaccine by inhibiting regulatory T cells"

### SUPPLEMENTAL MATERIALS

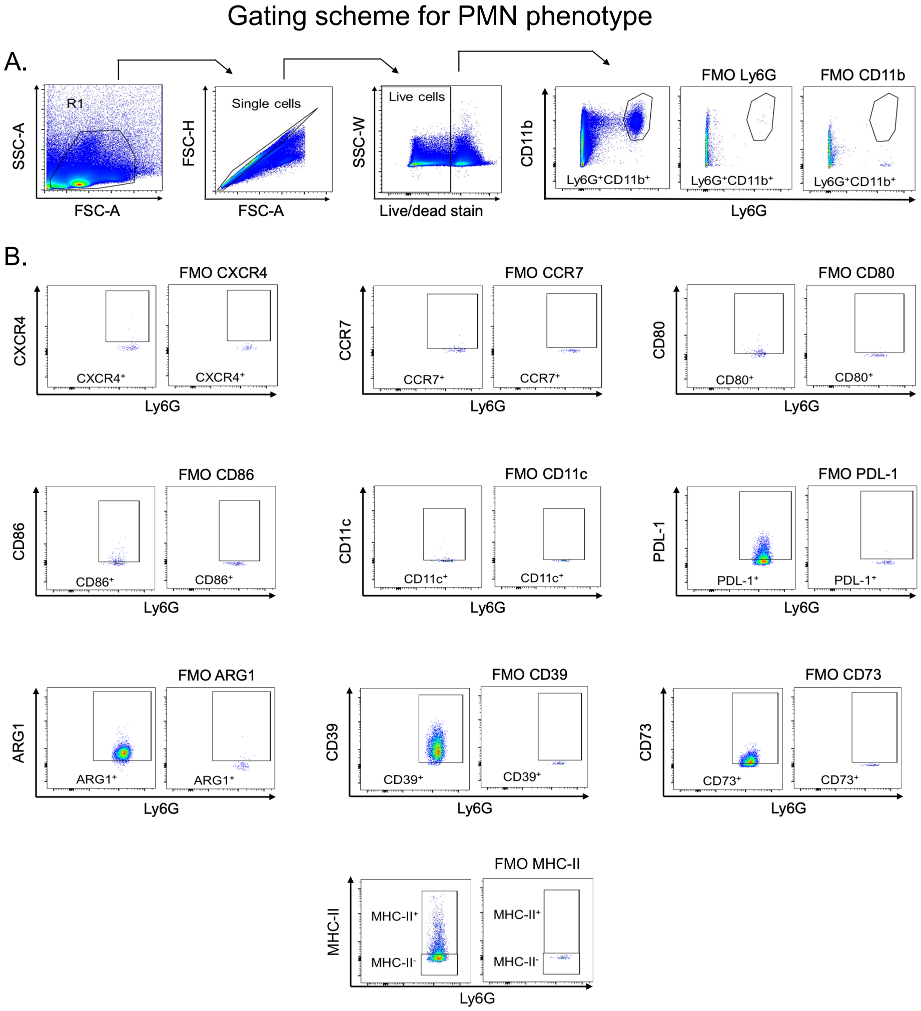


**Figure S1: Gating strategy for PMNs in mice**. Adult C57BL/6 female mice were injected with PCV or PBS. Blood, spleens, and vaccine-draining lymph nodes (vLNs) were collected from all mice and assessed for PMN phenotype by flow cytometry. The gating strategies are shown. (A) Live single cells were gated on and the percentage of PMNs (Ly6G$^+$CD11b$^+$) was determined. (B) The expression of CXCR4, CCR7, CD80, CD86, CD11c, PD-L1, ARG1, CD73, CD39 and MHC-II on PMNs were determined. Abbreviations: SSC-A = side scatter area; FSC-A = forward scatter area; FSC-H = forward scatter height; SSC-W = side scatter width; FMO = fluorescent minus one; MHC = Major Histocompatibility Complex; ARG= arginase.

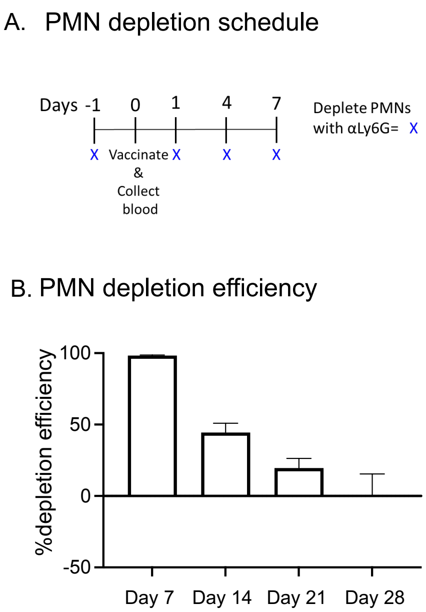


**Figure S2: PMN depletion efficiency.** Adult C57BL/6 female mice were injected with PCV or PBS and depleted of their PMNs using an αLy6G antibody or treated with an isotype control. (A) Timeline of PMNs depletion. (B) At the indicated times post vaccination, spleens were collected from all mice, and the depletion efficiency of PMNs was calculated at each timepoint using PMN numbers assessed by MPO ELISA. Data are pooled from n=7 mice per group at day 7 and n=4 mice per group at all other time points. Graph represents the mean +/- SD.

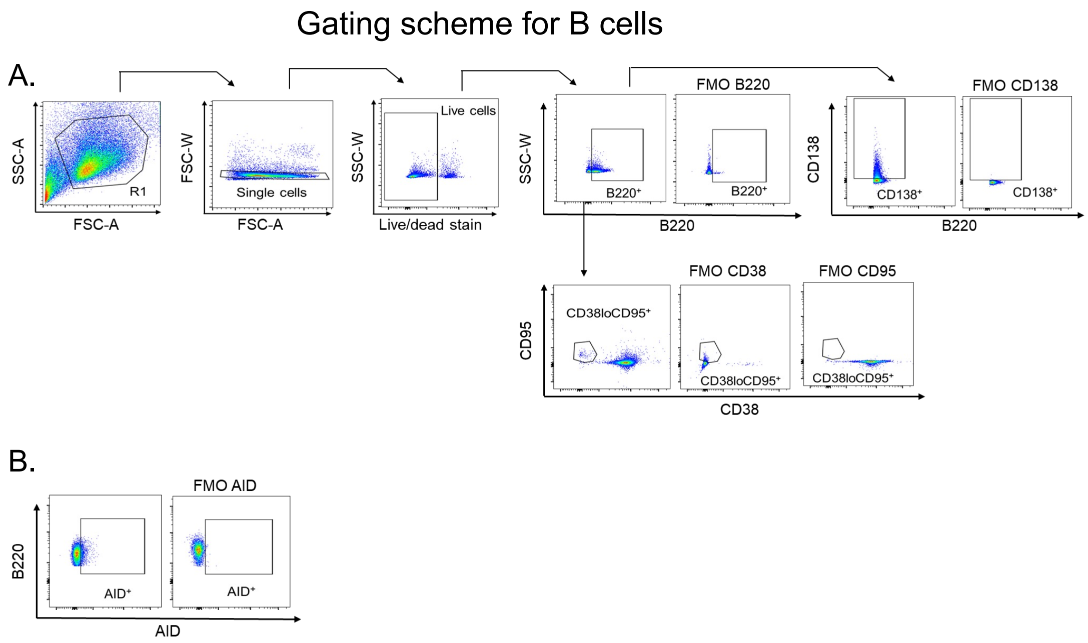


**Figure S3: Gating strategy for B cells and AID measurement.** Adult C57BL/6 female mice were injected with PCV or PBS and depleted of their PMNs or treated with an isotype control. Spleens were collected from all mice and assessed using flow cytometry. The gating strategies are shown. (A) Live single cells were gated on, and the percentages of total B cells (B220$^{+}$) followed by percentages of plasmablasts (CD138$^{+}$) and GC B cells (CD38$^{lo}$CD95$^{+}$) within the total B cell gate. (B) Percentage of Activation-induced Cytidine Deaminase (AID) expression by total B cells and GC B cells was determined. Abbreviations: SSC-A = side scatter area; FSC-A = forward scatter area; FSC-W = forward scatter width; SSC-W = side scatter width; FMO = fluorescent minus one.

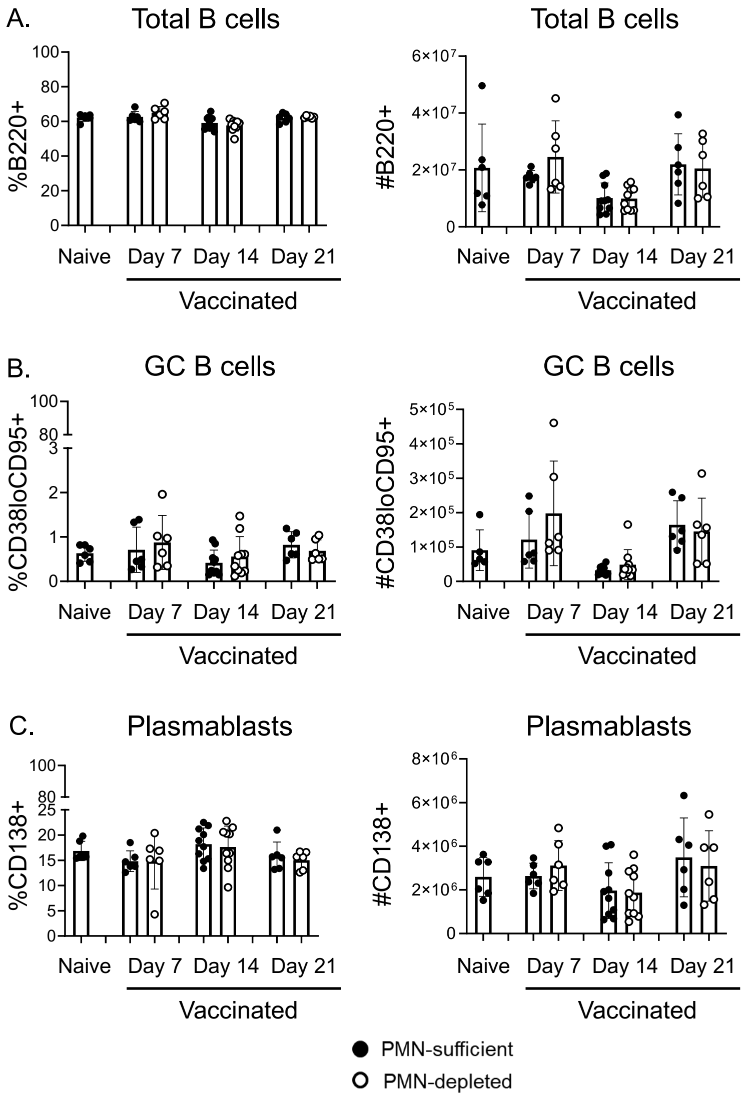


**Figure S4: Effect of PMN depletion on B cells in the spleen following PCV vaccination.** Adult C57BL/6 female mice were injected with PCV or PBS and depleted of their PMNs or treated with an isotype control. Spleens were collected from all mice and assessed using flow cytometry. Percentages (left) and numbers (right) of (A) total B cells (B220$^+$ from live gate), (B) Germinal center B cells (CD38$^{lo}$CD95$^+$ from B220$^+$ gate) and (C) plasmablasts (CD138$^+$ from B220$^+$ gate) were assessed at days 7, 14 and 21 post vaccination. Data are pooled from 4 separate experiments and each dot represents an individual mouse with a total of n=6 naïve, n=6 vaccinated day 7, n=10 vaccinated day 14, and n=6 vaccinated day 21 mice per group. Bar graphs represent the mean +/- SD.

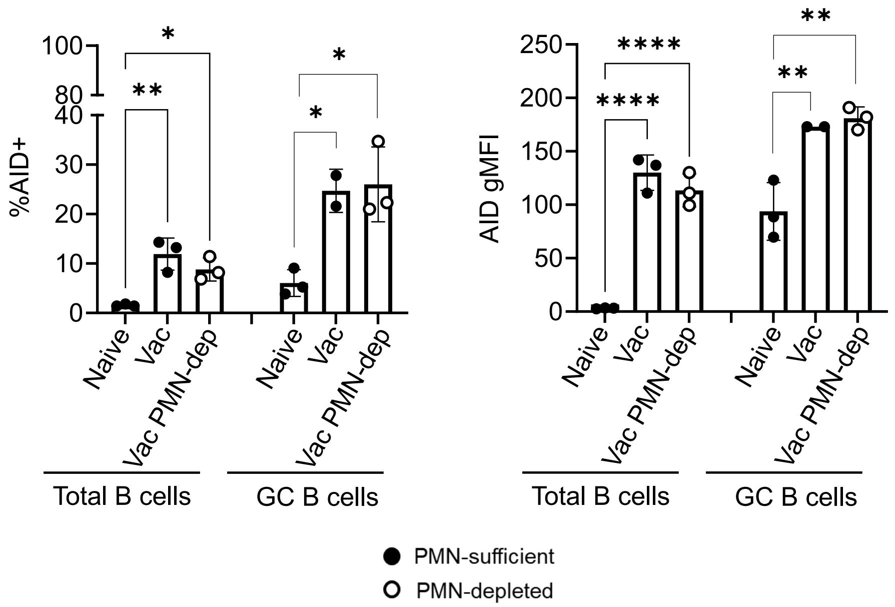


**Figure S5: Effect of PMN depletion on AID expression by B cells in the spleen following PCV vaccination.** Adult C57BL/6 female mice were injected with PCV or PBS and depleted of their PMNs or treated with an isotype control. Spleens were collected from all mice and assessed using flow cytometry. The percentages (left) and gMFI (right) expression of AID by total B cells (B220$^{+}$) and GC B cells (CD38$^{lo}$CD95$^{+}$) were assessed at 7 days post vaccination. Data are from 2 experiments and each dot represents an individual mouse with a total of n=2-3 mice per group. * denotes significant differences between the indicated groups as determined by One-Way ANOVA followed by Dunnett's multiple comparisons test. * is $p \leq 0.05$, ** is $p \leq 0.01$, and **** is $p \leq 0.0001$. Bar graphs represent the mean +/- SD.

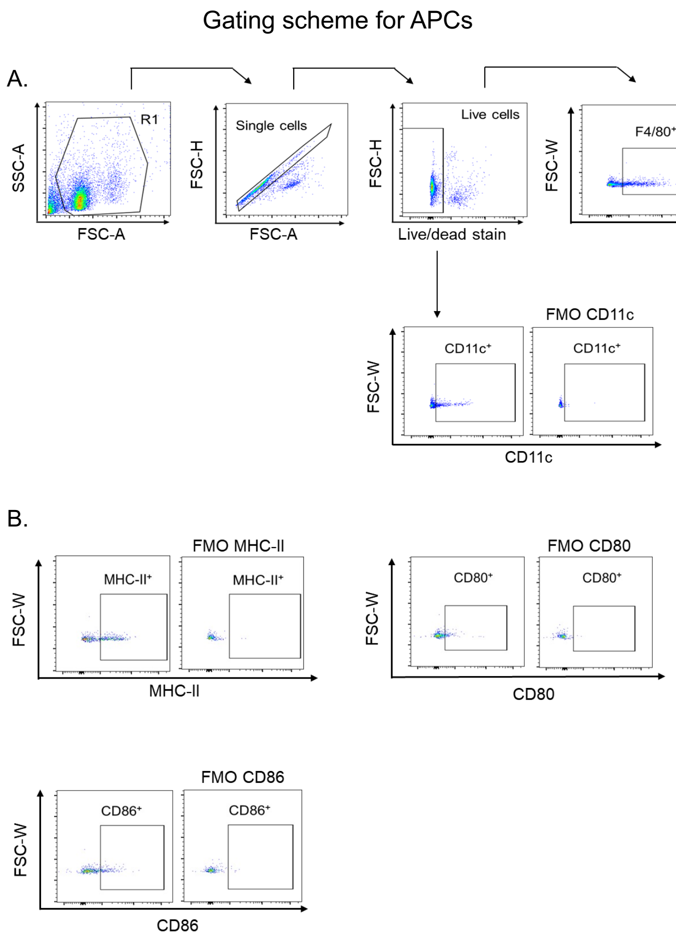


**Figure S6: Gating strategy for antigen presenting cells**. Adult C57BL/6 female mice were injected with PCV or PBS and depleted of their PMNs or treated with an isotype control. Spleens were collected from all mice and assessed for antigen presenting cells by flow cytometry. The gating strategies are shown. (A) The percentage of macrophages (F4/80$^+$) and dendritic cells (CD11c$^+$) were determined from live cells. (B) The percentage expression of CD80, CD86, and MHC-II CD11c on each cell type was determined. Abbreviations: SSC-A = side scatter area; FSC-A = forward scatter area; FSC-H = forward scatter height; FSC-W = forward scatter width; FMO = fluorescent minus one; MHC = Major Histocompatibility Complex.

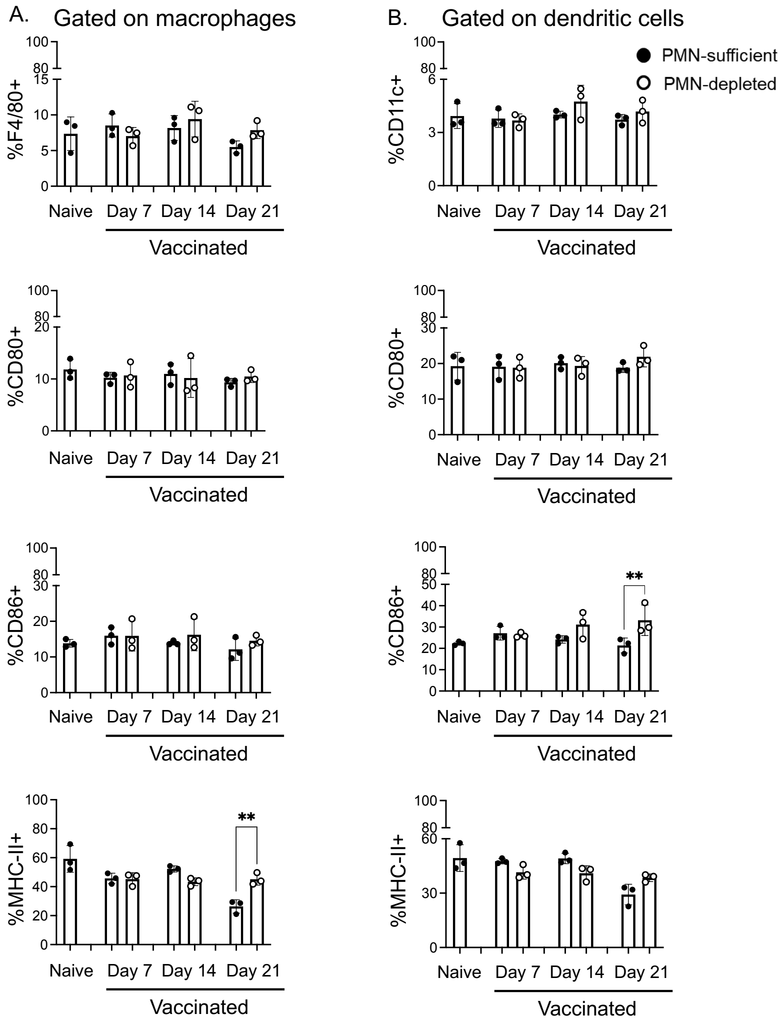


**Figure S7: Effect of PMN depletion on antigen presenting cells following PCV vaccination.** Adult C57BL/6 female mice were injected with PCV or PBS and depleted of their PMNs or treated with an isotype control. Spleens were collected from all mice and assessed using flow cytometry. The percentages of (A) macrophages (F4/80$^{+}$) and (B) dendritic cells (CD11c$^{+}$) in live cells were assessed at days 7, 14 and 21 post vaccination. Additionally, the percentage expression of CD80, CD86 and MHC-II on (A) macrophages and (B) dendritic cells was determined. Data are from 2 experiments and each dot represents an individual mouse with a total of n=3 mice per group. * denotes significant differences between the indicated groups as determined by One-Way ANOVA followed by Šídák's multiple comparisons test. ** is $p \leq 0.01$. Bar graphs represent the mean +/- SD. Abbreviations: MHC = Major Histocompatibility Complex.

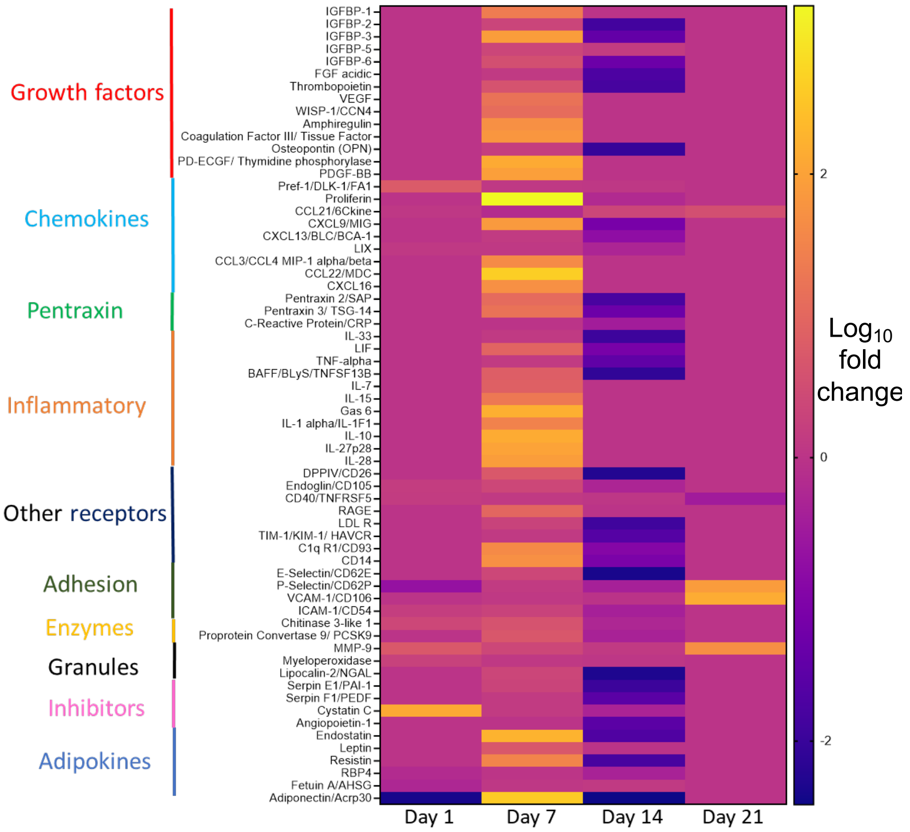


**Figure S8: Effect of PMNs on the inflammatory environment in the spleen following PCV vaccination.** Adult C57BL/6 female mice were injected with PCV and depleted of their PMNs or treated with an isotype control. Spleens were collected from all mice at days 1-, 7-, 14- and 21 post vaccination and assessed for cytokine/chemokines. Data are pooled from 3 separate experiments with 3 mice per group per timepoint and are presented in a heat map as $Log_{10}$ of the ratio of values from PMN-depleted mice divided by values from PMN-sufficient mice.

Gating scheme for IFNγ expression by PMNs

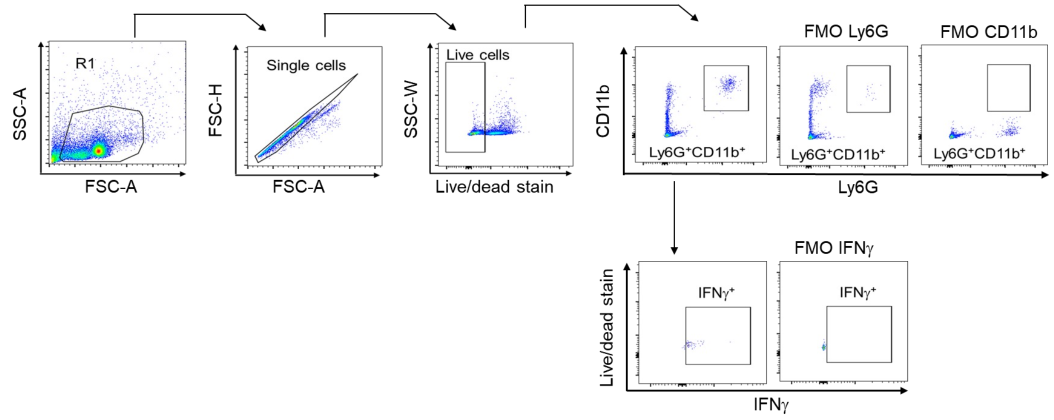


**Figure S9: Gating strategy for IFNγ expression by PMNs.** Adult C57BL/6 female mice were injected with PCV or PBS. Spleens were collected from all mice and assessed for IFNγ production by PMNs 18 hours post vaccination using flow cytometry. The gating strategy is shown. Live single PMNs (Ly6G$^{+}$CD11b$^{+}$) were gated on and percentage expression of IFNγ was determined. Abbreviations: SSC-A = side scatter area; FSC-A = forward scatter area; FSC-H= forward scatter height; SSC-W = side scatter width; FMO = fluorescent minus one.

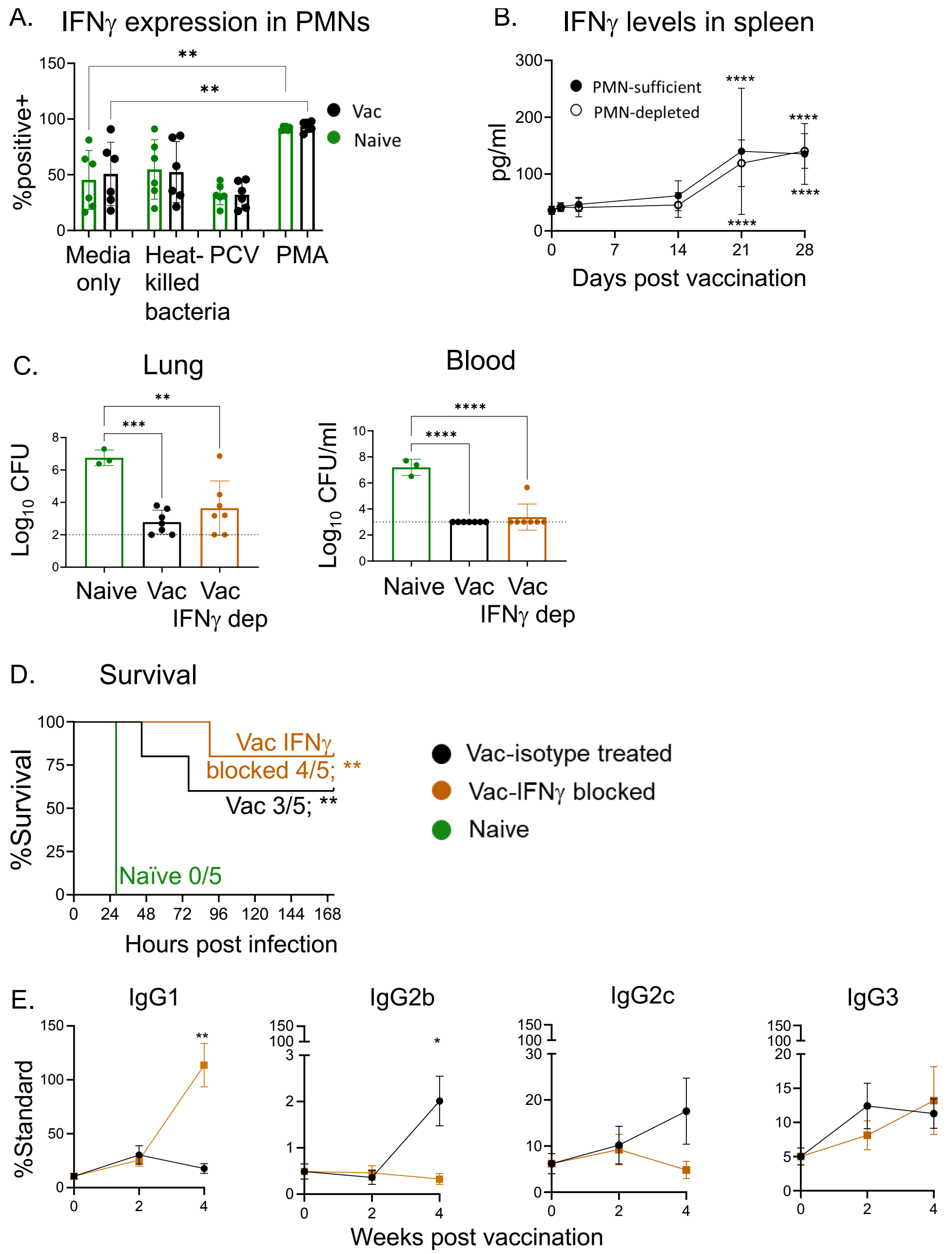


**Figure S10: Role of IFNγ in antibody responses to PCV.** (A) Adult C57BL/6 female mice were injected with PCV or PBS. Spleens were collected from all mice, stimulated with heat-killed *S. pneumoniae*, PCV, PMA or media alone, and assessed for IFNγ production by PMNs 18 hours post vaccination using flow cytometry. (B) Adult C57BL/6 female mice were injected with PCV or PBS and depleted of their PMNs or treated with an isotype control. Spleens were collected from mice across time following vaccination and assessed for IFNγ levels via ELISA. (C-E) Adult

C57BL/6 female mice were injected with PCV or PBS and treated with αIFNγ or an isotype control. All mice were infected 4 weeks following vaccination with $10^7$ CFU of *S. pneumoniae* TIGR4, assessed for antibody production, bacterial burden, and survival. (C) Lung and blood bacterial burdens were assessed 24 hours post infection. (D) Mice were monitored for survival for 7 days following infection. (E) Production of IgG subtypes was monitored on weeks 0, 2 and 4 post vaccination. (A) Data are pooled from 2 separate experiments and each dot represents an individual mouse with a total of n=6 mice per group. * denotes significant differences between the indicated groups as determined by One-Way ANOVA followed by Šídák's multiple comparisons test. (B) Data are pooled from separate experiments with at least n=3 mice per timepoint per group. Line graphs represent the mean +/- 95% confidence interval. * denotes significant differences from baseline as determined by One-Way ANOVA followed by Dunnett's multiple comparisons test. (C) Data are pooled from 2 separate experiments and each dot represents an individual mouse with a total of n=3 naïve, n=6 vaccinated isotype treated and n=6 vaccinated IFNγ-treated mice. * denotes significant differences between the indicated groups as determined by One-Way ANOVA followed by Tukey's multiple comparisons test. (D) Data are pooled from 2 separate experiments with a total of n=5 mice per group. * denotes significant differences from naïve as determined by Mantel-Cox test. (E) Data are pooled from separate experiments with n=11 mice per timepoint. Line graphs represent the mean +/- SEM. * denotes significant differences from baseline as determined by Kruskal-Wallis test followed by Dunn's multiple comparisons. * is $p \leq 0.05$, ** is $p \leq 0.01$, *** is $p \leq 0.001$, and **** is $p \leq 0.0001$.

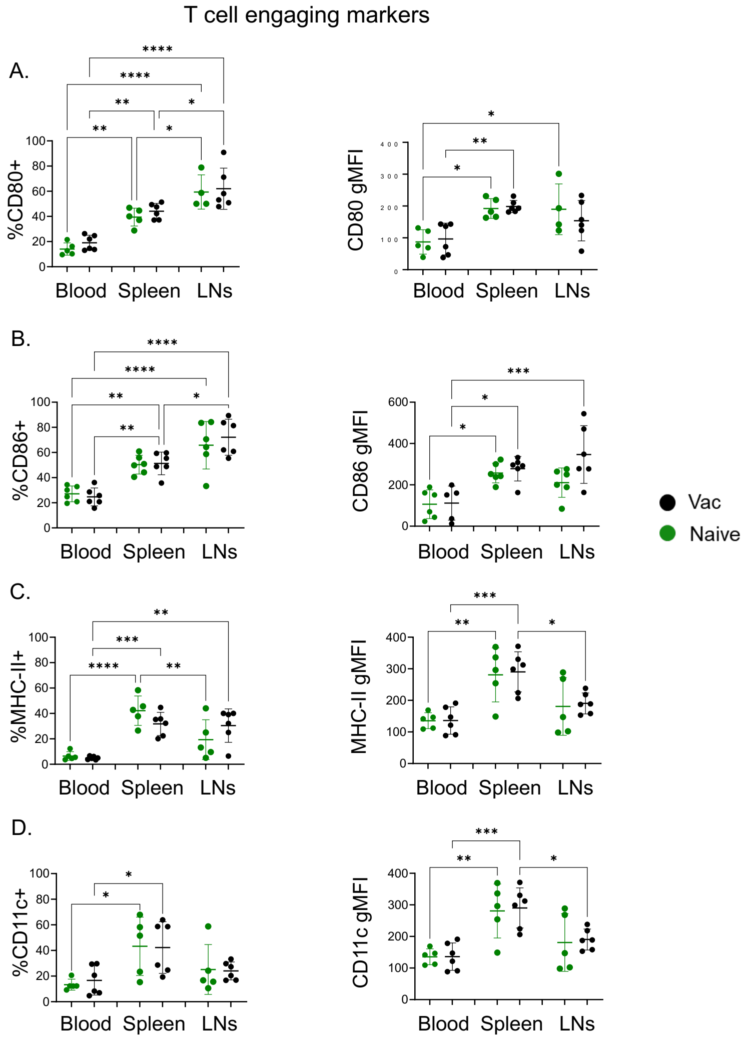


**Figure S11: PMN expression of T cell engaging markers following PCV vaccination.** Adult C57BL/6 female mice were injected with PCV or PBS. At 18- 24 hours post vaccination, blood, spleen and vLNs were collected and assessed using flow cytometry. Percentages (left) and gMFI (right) of (A) CD80, (B) CD86, (C) MHC-II, and (D) CD11c on live PMNs ($Ly6G^{+}CD11b^{+}$) were determined. Data are pooled from 3 separate experiments and each dot represents an individual mouse with a total of n=5 naïve and n=6 vaccinated mice. * denotes significant differences between the indicated groups as determined by One-Way ANOVA followed by Šídák's multiple comparisons test. * is $p≤0.05$, ** is $p≤0.01$, *** is $p≤0.001$, and **** is $p≤0.0001$. Graphs represent the mean +/- SD. Abbreviations: gMFI = geometric Mean Fluorescence Intensity; MHC = Major Histocompatibility Complex.

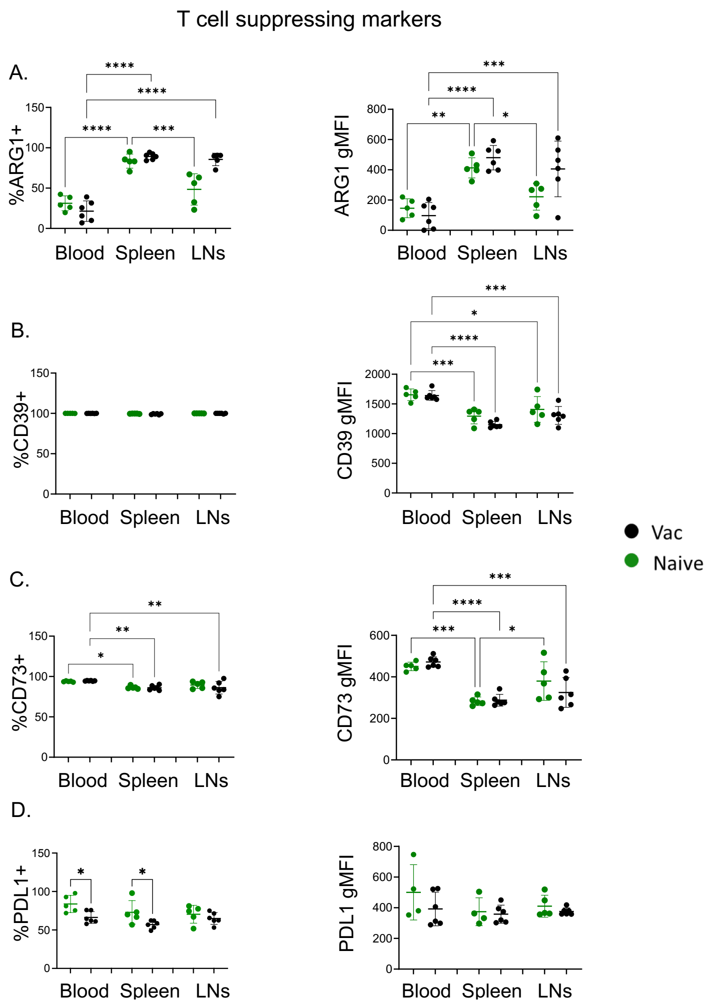


**Figure S12: PMN expression of T cell suppressing markers following PCV vaccination.** Adult C57BL/6 female mice were injected with PCV or PBS. At 18 hours post vaccination, blood, spleen and vLNs were collected and assessed using flow cytometry. Percentages (left) and gMFI (right) of (A) ARG1, (B) CD39, (C) CD73, and (D) PD-L1 on live PMNs (Ly6G$^+$CD11b$^+$) were determined. Data are pooled from 3 separate experiments and each dot represents an individual mouse with a total of n=5 naïve and n=6 vaccinated mice. * denotes significant differences between the indicated groups as determined by One-Way ANOVA followed by Šídák's multiple comparisons test. * and # is $p \leq 0.05$, ** is $p \leq 0.01$, *** is $p \leq 0.001$, and **** is $p \leq 0.0001$. Graphs represent the mean +/- SD. Abbreviations: gMFI = geometric Mean Fluorescence Intensity; ARG= arginase.

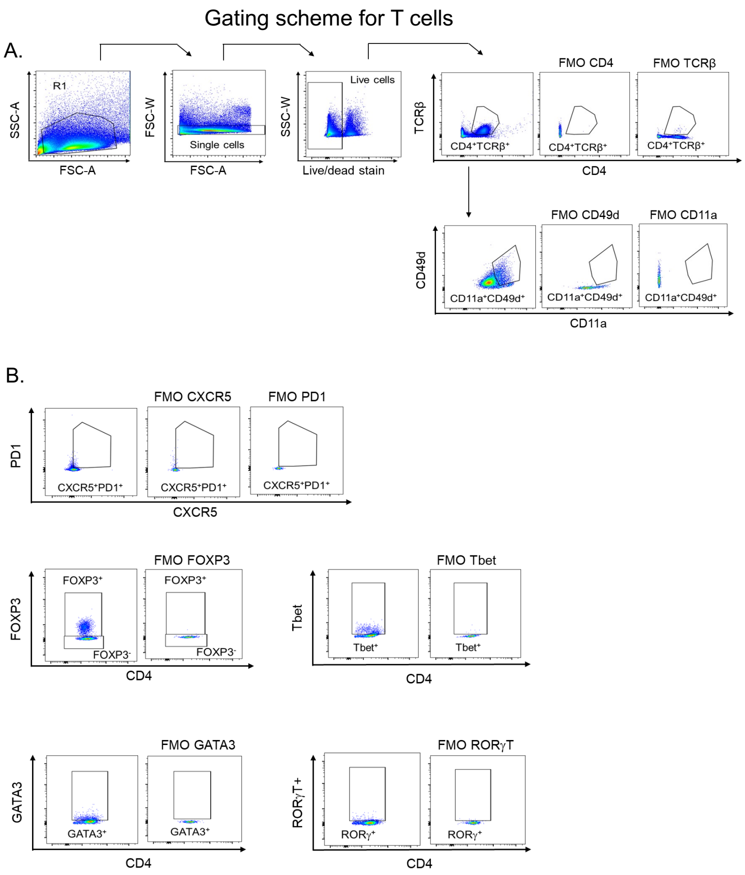


**Figure S13: Gating strategy for T cell subsets**. Adult C57BL/6 female mice were injected with PCV or PBS and depleted of their PMNs or treated with an isotype control. Spleens were collected from all mice and assessed using flow cytometry. The gating strategies are shown. (A) Live single cells were gated on and the percentages of total CD4 T cells ($CD4^{+}TCR\beta^{+}$) and antigen experienced ($CD11a^{+}CD49d^{+}$) T cells were determined. (B) Total and antigen experienced CD4 T cells were gated on and the percentages of T follicular helper cells ($CXCR5^{+}PD1^{+}$), Regulatory T cells ($FOXP3^{+}$), T helper 1 ($Tbet^{+}FOXP3^{-}$), Th2 ($GATA3^{+}FOXP3^{-}$) and Th17 ($ROR\gamma T^{+}FOXP3^{-}$) subsets were determined in each gate. Additionally, total and antigen experienced Tfhs were gated on and the percentage of T follicular regulatory cells ($CXCR5^{+}PD1^{+}$ $FOXP3^{+}$) cells were determined in each gate. Abbreviations: SSC-A = side scatter area; FSC-A = forward scatter area; FSC-W = forward scatter width; SSC-W = side scatter width; FMO = fluorescent minus one; TCR = T cell receptor.

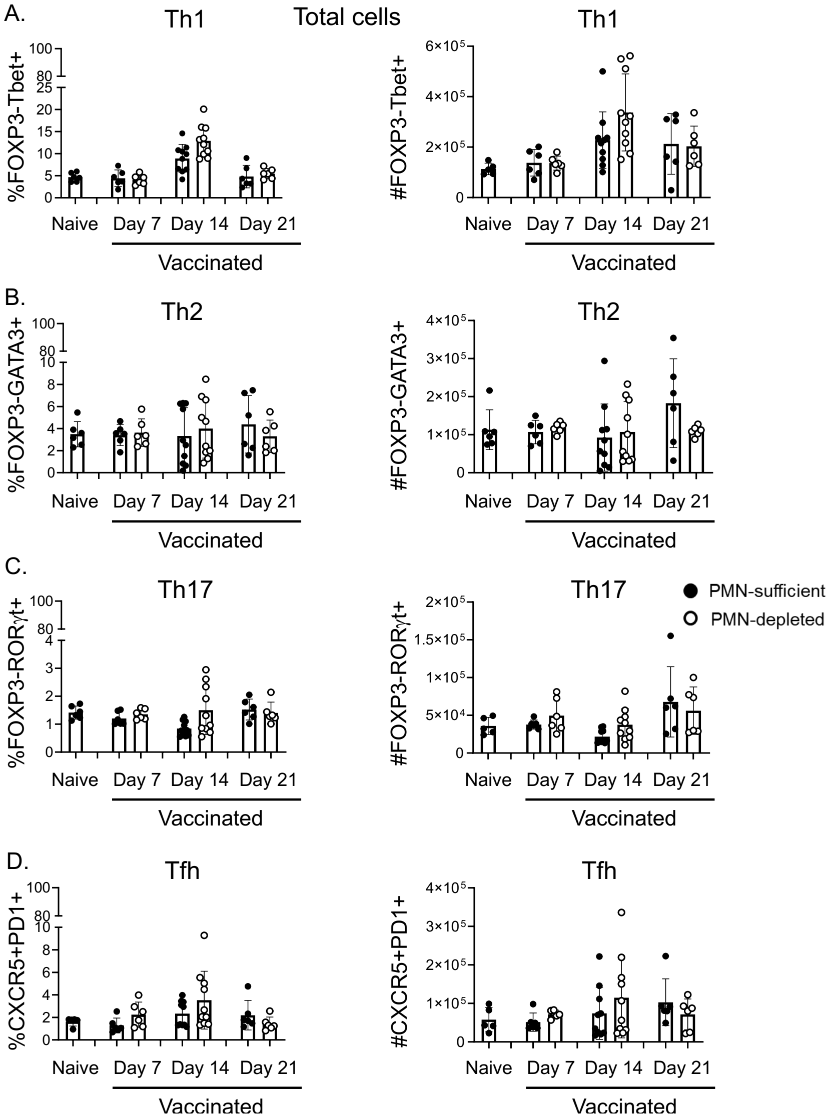


**Figure S14: Effect of PMN depletion on total T helper cell subsets.** Adult C57BL/6 female mice were injected with PCV or PBS and depleted of their PMNs or treated with an isotype control. Spleens were collected from all mice and assessed using flow cytometry. Total T cells ($CD4^+TCR\beta^+$) were gated on and percentages (left) and numbers (right) of different T cell subsets were determined in each gate at days 7, 14 and 21 post vaccination. (A) T helper 1 ($Tbet^+FOXP3^-$), (B) Th2 ($GATA3^+FOXP3^-$), (C) Th17 ($ROR\gamma T^+FOXP3^-$), and (D) T follicular helper cell ($CXCR5^+PD1^+$) were gated on. Data are pooled from 4 separate experiments and each dot represents an individual mouse with a total of n=6 naïve, n=6 vaccinated day 7, n=10 vaccinated day 14, and n=6 vaccinated day 21 mice per group. Bar graphs represent the mean +/- SD.

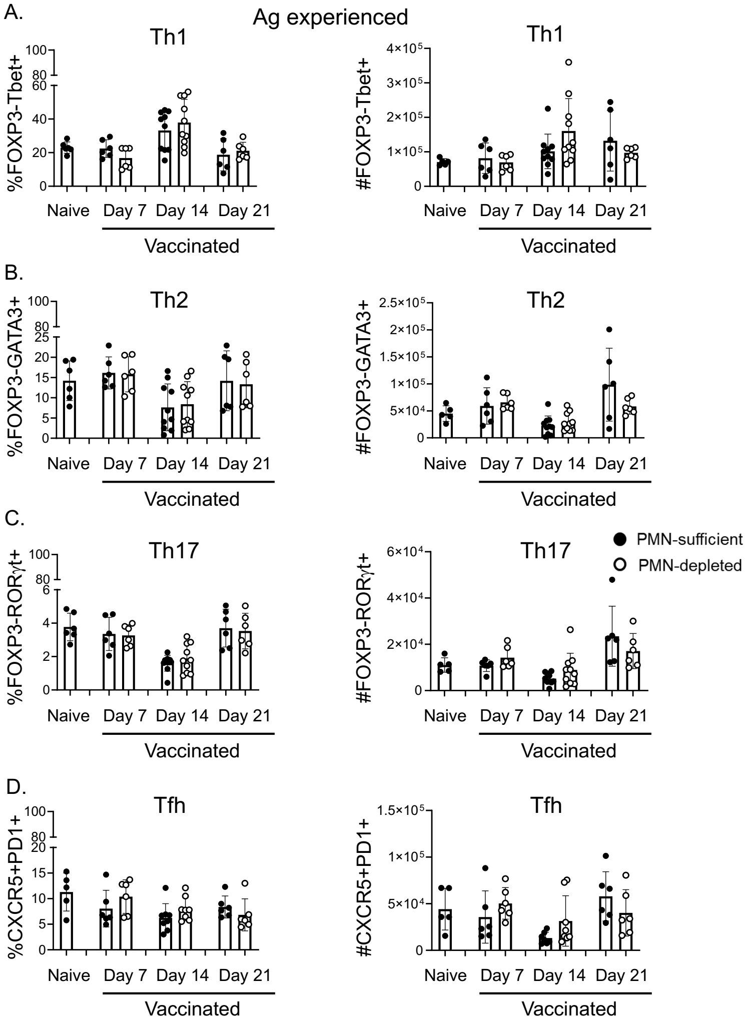


**Figure S15: Effect of PMN depletion on antigen experienced T helper cell subsets.** Adult C57BL/6 female mice were injected with PCV or PBS and depleted of their PMNs or treated with an isotype control. Spleens were collected from all mice and assessed using flow cytometry. Antigen experienced T cells (CD4$^+$TCRβ$^+$ CD11a$^+$CD49d$^+$) were gated on and percentages (left) and numbers (right) of different T cell subsets were determined in each gate at days 7, 14 and 21 post vaccination. (A) T helper 1 (Tbet$^+$FOXP3$^-$), (B) Th2 (GATA3$^+$FOXP3$^-$), (C) Th17 (RORγT$^+$FOXP3$^-$), and (D) T follicular helper cell (CXCR5$^+$PD1$^+$) were gated on. Data are pooled from 4 separate experiments and each dot represents an individual mouse with a total of n=6 naïve, n=6 vaccinated day 7, n=10 vaccinated day 14, and n=6 vaccinated day 21 mice per group. Bar graphs represent the mean +/- SD.

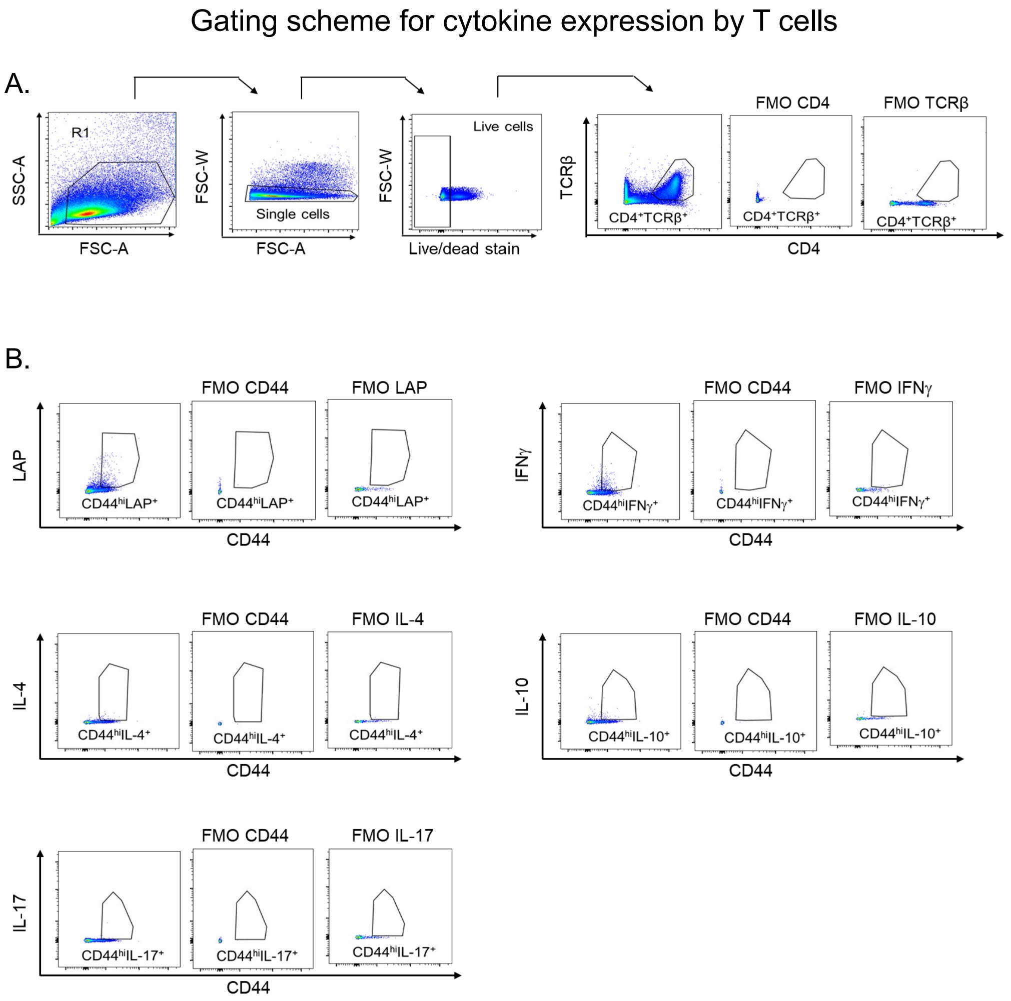


**Figure S16: Gating strategy for cytokine expression by T cells.** Adult C57BL/6 female mice were injected with PCV or PBS and depleted of their PMNs or treated with an isotype control. On day 14 post vaccination, spleens were collected from all mice and assessed for cytokine production by T cells using flow cytometry. The gating strategies are shown. (A) Live single T cells ($CD4^{+}TCR\beta^{+}$) were gated on and (B) percentage expression of IL-4, IL-10, IFNγ, LAP and IL-17 was determined. Abbreviations: SSC-A = side scatter area; FSC-A = forward scatter area; FSC-W = forward scatter width; FMO = fluorescent minus one; TCR = T cell receptor; LAP= latency-associated peptide.

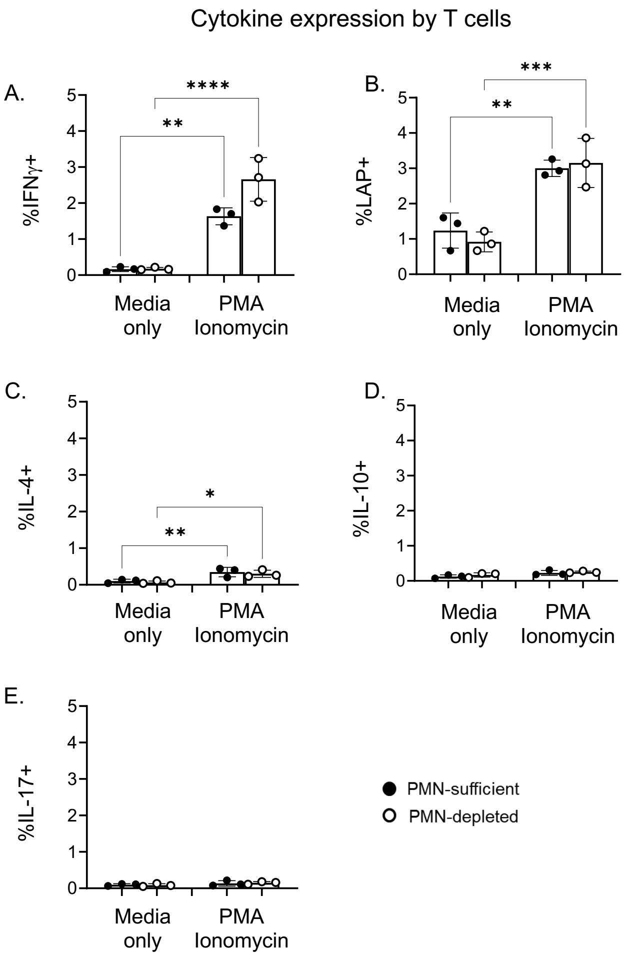


**Figure S17: Effect of PMNs on cytokine expression by T cells.** Adult C57BL/6 female mice were injected with PCV or PBS and depleted of their PMNs or treated with an isotype control. On day 14 post vaccination, spleens were collected from all mice, stimulated with PMA/Ionomycin or media alone, and assessed for cytokine production by T cells using flow cytometry. T cells ($CD4^{+}TCR\beta^{+}$) were gated on and percentage expression of IL-4, IL-10, IFN$\gamma$, LAP and IL-17 was determined. Data are from 2 experiments and each dot represents an individual mouse with a total of n=3 mice per group. * denotes significant differences between the indicated groups as determined by One-Way ANOVA followed by Šídák's multiple comparisons test. * is $p \leq 0.05$, ** is $p \leq 0.01$, *** is $p \leq 0.001$, and **** is $p \leq 0.0001$. Abbreviations: LAP= latency-associated peptide.

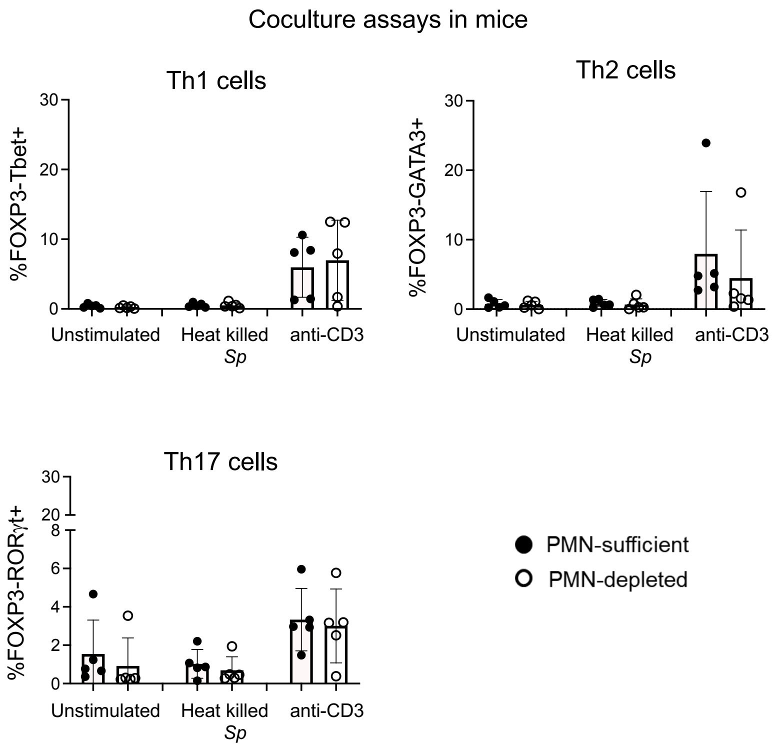


**Figure S18: Effect of PMN depletion on T helper cell subsets in mice *in vitro.*** Adult C57BL/6 female mice were injected with PCV or PBS and depleted of their PMNs or treated with an isotype control. On day 1 post vaccination, splenocytes were isolated and cultured in the presence or absence of PMNs for 2 days with heat-killed *S. pneumoniae* or αCD3 as stimulants. Percentage of (A) Th1, (B) Th2 and (C) Th17 cells in the CD4$^{+}$ gate are shown. Data are pooled from separate experiments.

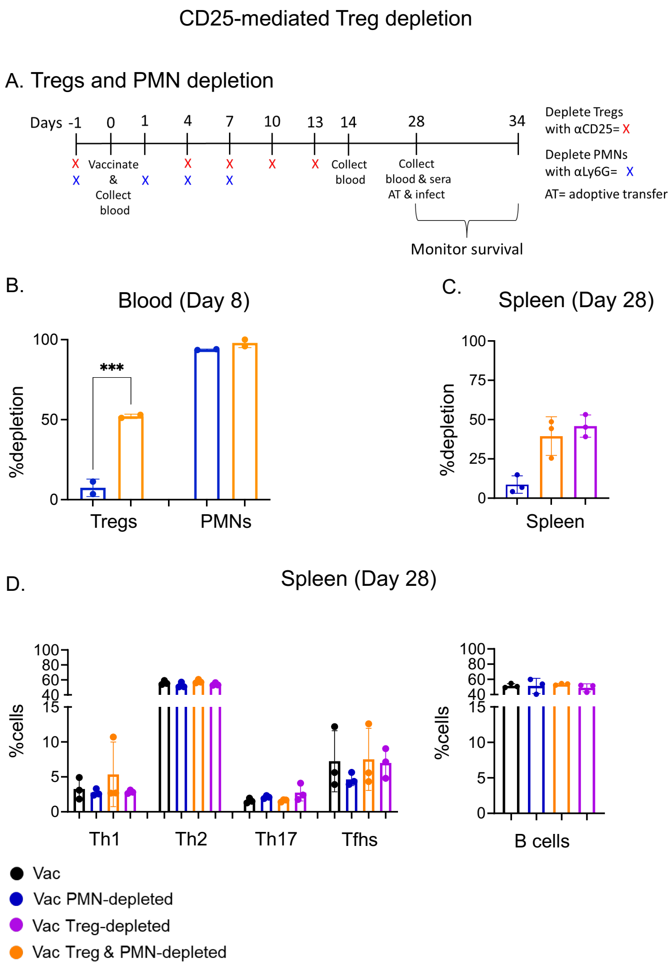


**Figure S19: Depletion of Tregs using an αCD25 depleting antibodies.** (A) Adult C57BL/6 female mice were injected with PCV or PBS and depleted of their PMNs and/or Tregs or treated with isotype controls. (B) On day 8 post vaccination, blood was collected from all mice and the depletion efficiencies of PMNs and Tregs were assessed using flow cytometry. (C&D) On day 28 post vaccination, spleens were collected from all mice and assessed for B and T cells. (C) Treg depletion efficiency and (D) the percentages of Th1, Th2, Th17, Tfhs and B cells were assessed. (B) Data show two technical replicates from two experiments, with samples pooled from n=8 mice per replicate. *** is $p \leq 0.001$ and denotes significant differences between the indicated groups as determined by One-way ANOVA followed by Šídák's multiple comparisons test. (C&D) Data are pooled from n=3 mice per group per timepoint. Bar graphs represent the mean +/- SD.

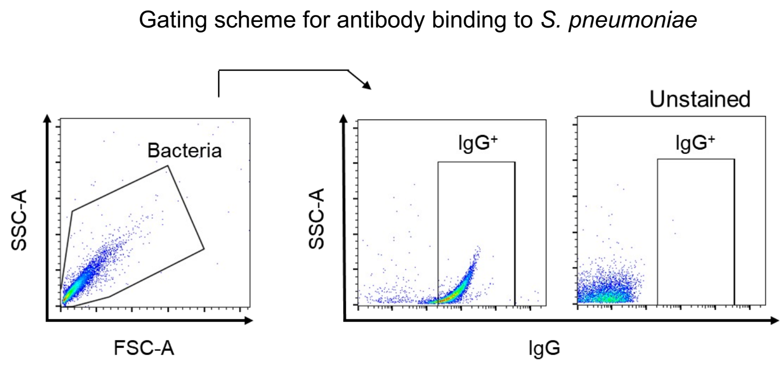


**Figure S20: Gating strategy for antibody binding to the surface of *S. pneumoniae*.** Adult C57BL/6 female and DEREG male mice were injected with PCV or PBS and depleted of their PMNs and/or Tregs or treated with an isotype/vehicle control. Sera collected 4-weeks post vaccination were used to assess the binding efficacy of antibodies to *S. pneumoniae* via flow cytometry. Antibody bind to the bacteria were detected with an APC-tagged anti-mouse IgG. Bacteria were gated on and the percentage of $IgG^+$ bacteria were determined. Abbreviations: SSC-A = side scatter area; FSC-A = forward scatter area.

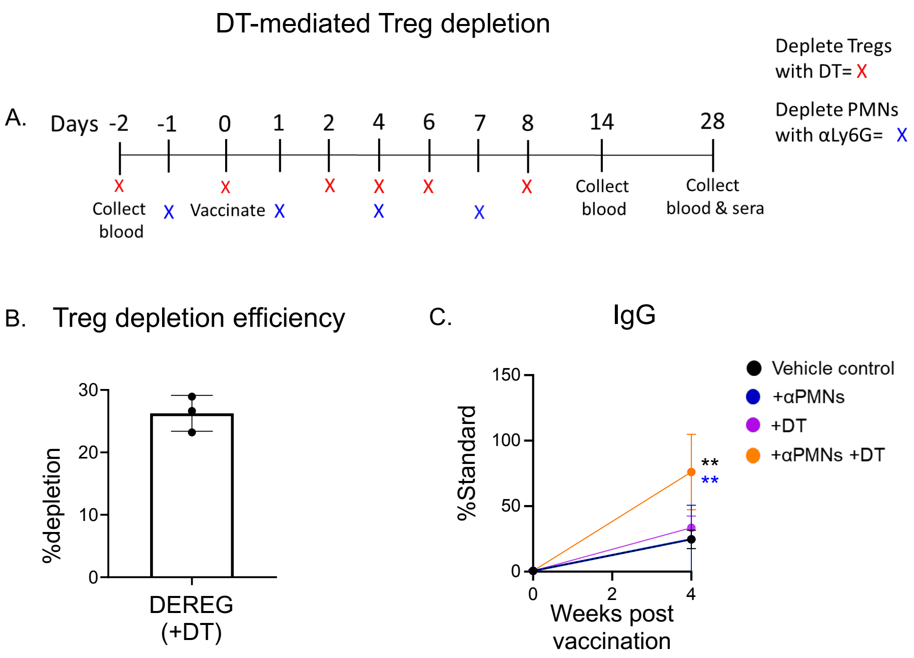


**Figure S21: Depletion of Tregs using diphtheria toxin in DEREG mice.** Adult DEREG male mice were injected with PCV or PBS and depleted of their PMNs and/or Tregs or treated with a vehicle control. (A) Timeline schedules for depletions is shown. (B) On day 14 post vaccination, spleens were collected from all mice and the depletion efficiency of Tregs was assessed using flow cytometry. Sera collected 4-weeks post vaccination were used to assess (C) total IgG antibodies against type 4 pneumococcal polysaccharides. (C) Data are presented as percentage of standard sera and pooled from n=4 mice per group. * denotes significant differences between PMNs- & Tregs-depleted mice and group matching the symbol color as determined by One-way ANOVA followed by Šídák's multiple comparisons test. All line graphs represent the mean +/- 95% confidence interval and all bar graphs represent the mean +/- SD.

Gating scheme for PMN phenotype in human donors

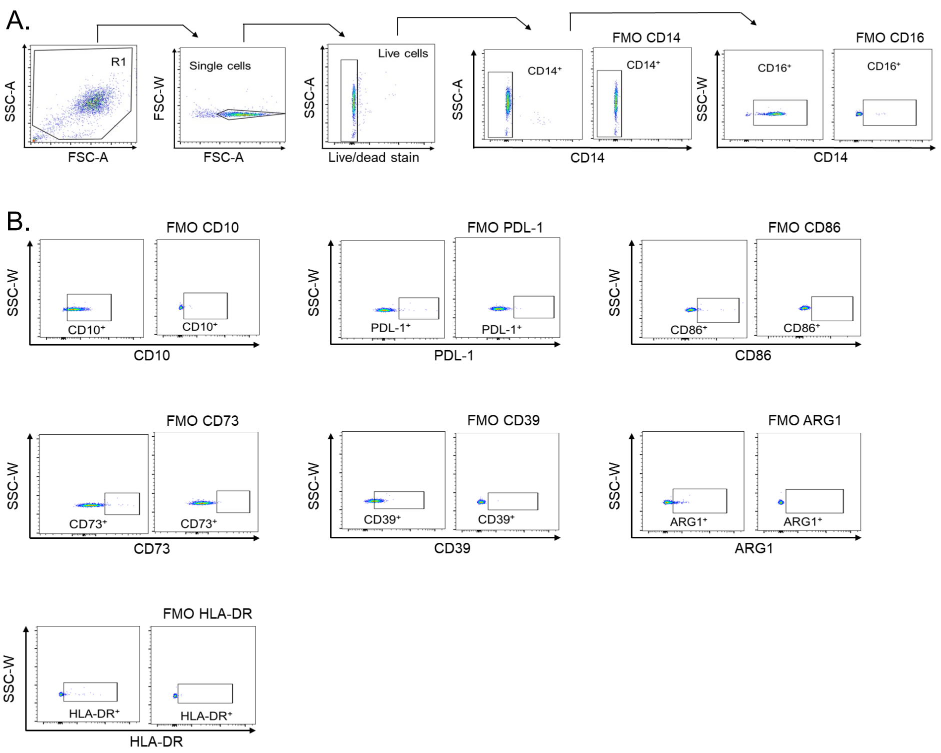


**Figure S22: Gating strategy for PMN phenotype in humans**. Young (24-29 years old) female donors were injected with PCV on day 0. Blood was collected prior to and on weeks 1 & 4 post vaccination. PMNs were isolated from their peripheral blood and assessed using flow cytometry. (A) CD16$^+$ cells were gated from CD14$^-$ live single cells. (B) CD16$^+$ cells were gated on and the percentages HLA-DR, CD10, CD86, PD-L1, ARG1, CD73 and CD39 were determined. Abbreviations: SSC-A = side scatter area; FSC-A = forward scatter area; FSC-W = forward scatter width; SSC-W = side scatter width; FMO = fluorescent minus one.

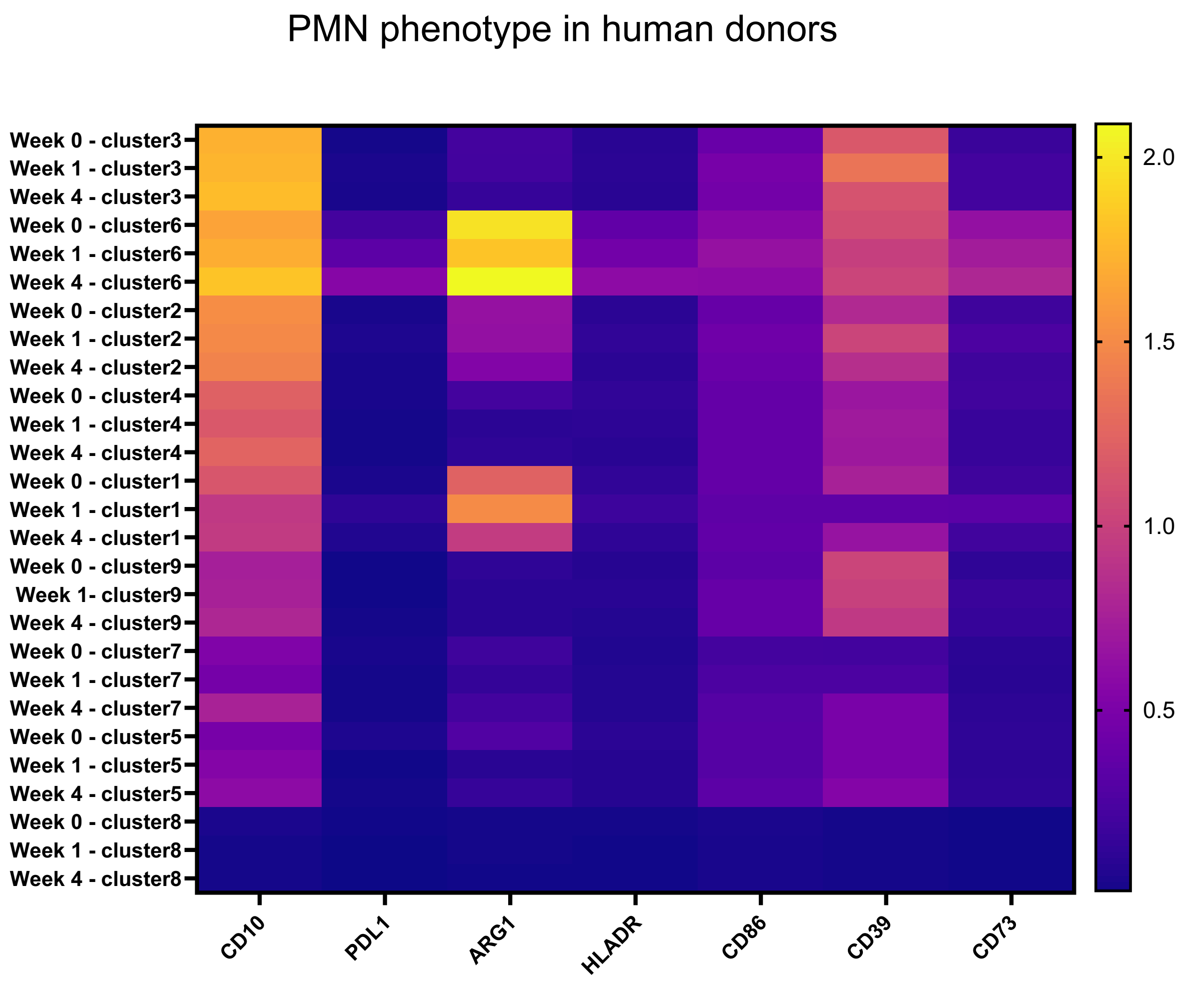


**Figure S23: Heat map of PMN phenotype in humans.** Young (24-29 years old) female donors were vaccinated with PCV. Blood was collected prior to (week 0) and on weeks 1 & 4 post vaccination. PMNs were isolated from their peripheral blood and assessed using flow cytometry. Expressions of HLA-DR, CD10, CD86, PD-L1, ARG1, CD73 and CD39 on PMNs were determined. Heat map of markers throughout the different weeks and clusters are shown. Data are pooled from six donors.

Gating scheme for T cells in coculture assays in human donors

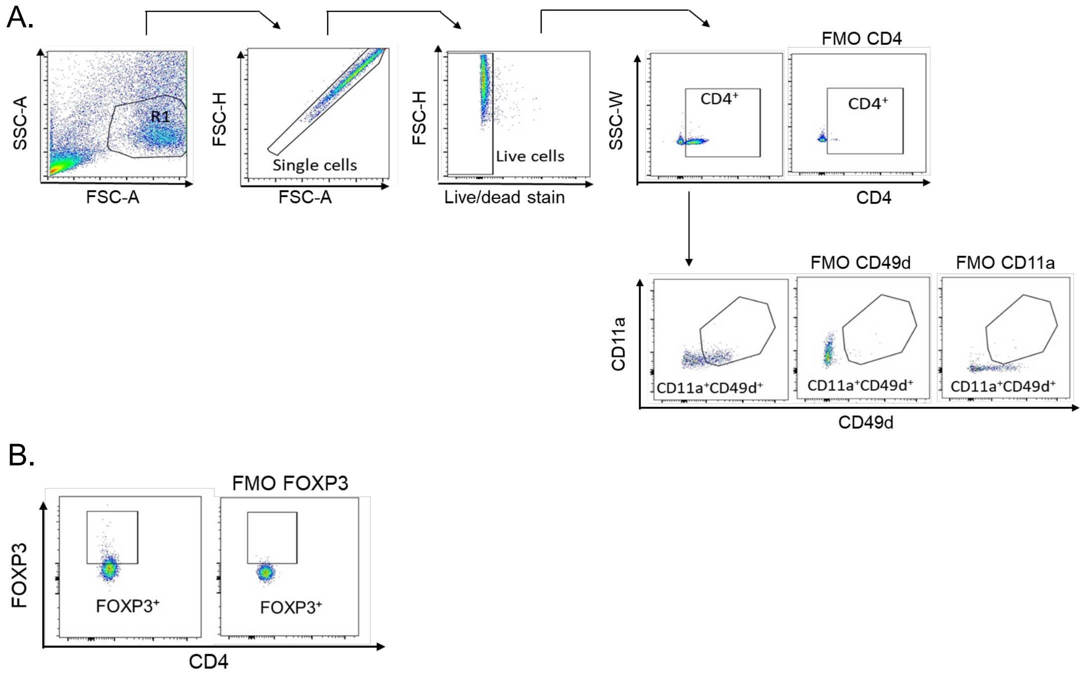


**Figure S24: Gating strategy for FOXP3$^+$ T cells in human coculture assays**. Young (24-29 years old) female donors were vaccinated with PCV. One week post vaccination, PBMCs and PMNs were isolated from their peripheral blood. PBMCs were cultured in the presence or absence of donor matched PMNs for 2 days with PCV as a stimulant. (A) Live single cells were gated on and the percentages of total CD4 T cells (CD4$^+$) and antigen experienced (CD11a$^+$CD49d$^+$) T cells were determined. (B) Total and antigen experienced CD4 T cells were gated on and the percentages of FOXP3$^+$ T cells were determined. Abbreviations: SSC-A = side scatter area; FSC-A = forward scatter area; FSC-H = forward scatter height; SSC-W = side scatter width; FMO = fluorescent minus one.
